# A single cell atlas reveals unanticipated cell type complexity in *Drosophila* ovaries

**DOI:** 10.1101/2021.01.21.427703

**Authors:** Maija Slaidina, Selena Gupta, Torsten Banisch, Ruth Lehmann

**Affiliations:** HHMI, Skirball Institute of Biomolecular Medicine, Department of Cell Biology, NYU Grossman School of Medicine, NY, USA, 540 First Avenue, NYC, NY 10016; Whitehead Institute and Department of Biology at MIT, 455 Mainstreet, Cambridge, MA 02142

## Abstract

Organ function relies on the spatial organization and functional coordination of numerous cell types. The *Drosophila* ovary is a widely used model system to study the cellular activities underlying organ function, including stem cell regulation, cell signaling and epithelial morphogenesis. However, the relative paucity of cell type specific reagents hinders investigation of molecular functions at the appropriate cellular resolution.

Here, we used single cell RNA sequencing to characterize all cell types of the stem cell compartment and early follicles of the *Drosophila* ovary. We computed transcriptional signatures and identified specific markers for nine states of germ cell differentiation, and 23 somatic cell types and subtypes. We uncovered an unanticipated diversity of escort cells, the somatic cells that directly interact with differentiating germline cysts. Three escort cell subtypes reside in discrete anatomical positions, and express distinct sets of secreted and transmembrane proteins, suggesting that diverse micro-environments support the progressive differentiation of germ cells. Finally, we identified 17 follicle cell subtypes, and characterized their transcriptional profiles. Altogether, we provide a comprehensive resource of gene expression, cell type specific markers, spatial coordinates and functional predictions for 34 ovarian cell types and subtypes.

## Introduction

Most organs are composed of numerous cell types. Their spatial organization determines the cellular interactions that ensure sustainable organ function, which lasts for the duration of an organism's life.

The fly ovary is an attractive and widely used model system to study the general principles of organ function, due to a combination of genetic tractability, small size, and cell type complexity. Oogenesis, or egg production, is the major function of the ovary. Adult ovaries comprise cells of two distinct lineages: the somatic cells of mesodermal origin, and the germ cells (GCs) that arise from primordial germ cells and differentiate into eggs. Ovaries are composed of 16 to 20 units called ovarioles, each of which serving as an egg production line (Figure 1C). At the anterior tip of an ovariole is a stem cell compartment, called the germarium, which houses germline stem cells (GSC), somatic follicle stem cells (FSCs) and their support cells. More posteriorly, follicles, or egg chambers, of increasingly advanced stages are lined up, with the mature oocytes located distally. Each follicle contains a germline cyst surrounded by a follicular epithelium, and will produce an egg. Specialized follicle cells, called stalk cells, separate follicles from one another, while polar cells at each pole of the follicle serve as organizers.

**Figure 1.**
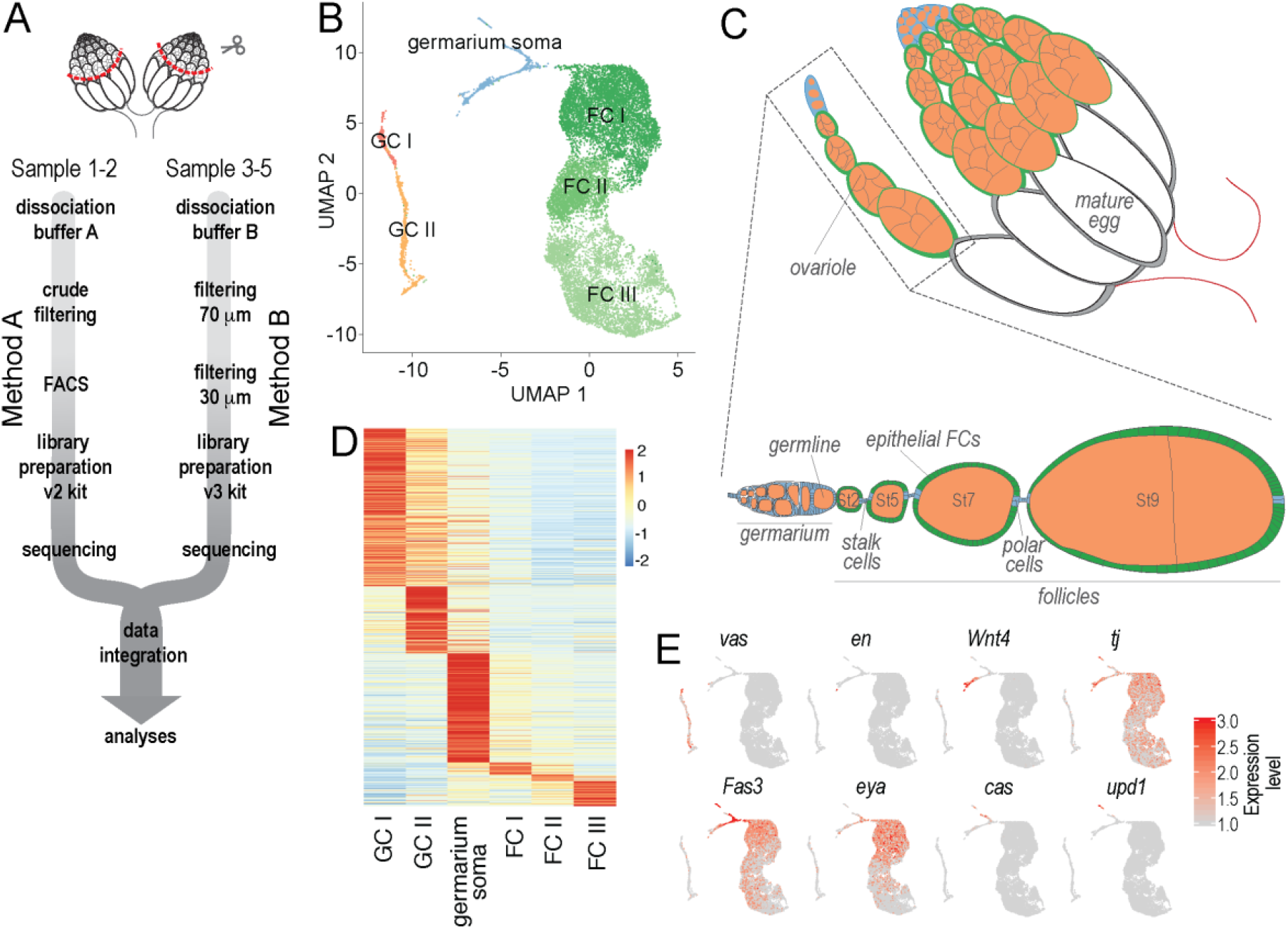
scRNA-seq of Drosophila adult ovaries. A - Schematic of the scRNA-seq sample preparation. B - UMAP plot of the entire dataset. Each dot represents a transcriptome of a single cell, and is color-coded according to cluster membership. C - Schematic drawing of an adult ovary. D - A heatmap visualizing average gene expression levels of all marker genes in each cluster. Red indicates highest and blue lowest expression. E - Expression of *vas, en, Wnt4, tj, Fas3, eya, cas* and *upd1*. Red indicates highest, and light grey lowest expression.

Both stem cell populations are excellent models to study stem cell maintenance and differentiation, and stem cell-niche interactions. GSC self-renewal has been extensively studied, and relies on signals from and adhesion to the niche, which comprise terminal filament (TF) and cap cells (CC) (Figure 3B). On the other hand, the mechanisms governing FSC self-renewal and differentiation are poorly understood.

Numerous somatic cell types support the multi-stage process of GC differentiation into mature eggs. Specifically, escort cells (ECs), also called inner germarium sheath cells, send protrusions around GCs and have a dual role: to promote both GSC renewal and GC differentiation (Kirilly et al. 2011, Wang and Page-McCaw 2018). This regulation relies on large surface contacts between the two cell types, and it is unclear whether distinct EC populations execute each function (Schulz et al. 2002, Banisch et al. 2017). Oogenesis is further supported by follicle cells (FCs) that arise from FSCs. FCs form an epithelial layer around the germ line-derived oocyte and nurse cells, thus delimiting the follicles. FCs are essential for egg development; they determine the body axes of the future embryo, and secrete yolk proteins and the eggshell. Numerous signaling pathways are involved in specification and morphogenesis of follicle cell subtypes. Therefore, the fly ovary also serves as a powerful experimental paradigm to dissect the molecular basis of cell-cell signaling. Despite the identification of various genes regulating oogenesis, the persistent paucity of cell type specific markers and tools has hindered progress in determining the exact role of these factors with the appropriate cellular resolution. For example, specific markers for GSCs, FSC and other somatic cell types remain to be identified.

To bridge this gap, we used a single-cell RNA sequencing (scRNA-seq) approach to systematically identify and characterize all cell types that compose the stem cell compartment and early-to-mid stage egg chambers. Stepwise clustering produced 34 cell clusters, allowing us to uncover new markers and determine precise cell type identities by analyzing *in situ* gene expression in ovaries. Our work revealed an unanticipated diversity of somatic cells in the germarium, including three subtypes of escort cells. Moreover, we identified 17 epithelial follicle cell subtypes, and characterized 9 distinct states of GC differentiation extending from earliest stages of differentiation to stage 2 of oogenesis. We computed transcriptional signatures, and tentatively predicted molecular functions for all ovarian cell types and subtypes. The novel markers will enable future tool development and functional analyses. Finally, we integrate our findings with recently published ovary atlases, and discuss the novel insights for the field of reproductive biology in *Drosophila*.

## Results

### scRNA-seq using two distinct methods produce equivalent results

We aimed to obtain and characterize single cell transcriptomes from the germarium and early follicles to identify cell type specific markers, and thus enable further functional studies. We used a nanoliter droplet based 10x Genomics Chromium system for single cell RNA sequencing (Figure 1A). We performed the experiment in five replicates and selected high quality transcriptomes (Supplemental Text). Despite using two different methods for cell dissociation, cell suspension preparation, library preparation, and sequencing depth, all five sequencing experiments produced similar results (Supplemental Text, Figure 1A, Supplemental Figure S1A-C). Data from all replicates were aligned and merged using the Seurat 3 algorithm for library integration (Figure S1A).

Our initial coarse clustering revealed six clusters, visualized on a UMAP (Uniform Manifold Approximation and Projection) plot, where each dot represents a single cell transcriptome (Figure 1B). Two clusters expressed the GC marker *vas* (Lasko and Ashburner 1988), and were thus annotated as germ cells (clusters GC I and II) (Figure 1B,1E). The three largest clusters expressed the follicle cell markers *Fas3* and *eya* (Nystul and Spradling 2007, Bai and Montell 2002), and therefore, corresponded to epithelial follicle cells (clusters FC I, II, and III) (Figure 1B, 1E). Finally, the last cluster contained somatic cell types of the germarium - TF and CC that express *en*, ECs that express *Wnt4* and *tj* (Forbes et al. 1996, Mottier-Pavie et al. 2016, Li et al. 2003, Kawashima et al. 2003), early follicle cell lineages that express *Fas3*, *tj* and *eya* (Nystul and Spradling 2007, Li et al. 2003, Kawashima et al. 2003, Bai and Montell 2002), and stalk and polar cell lineages that express *cas* (Chang et al. 2013) and *upd1* (Silver and Montell 2001) (cluster germarium soma) (Figure 1B, 1E). We did not detect a separate cluster containing the muscle cells that ensheath the ovarioles (epithelial sheath cells) or peritoneal sheath, suggesting that those cells were lost during dissections or dissociations.

To compare the transcriptional signatures of these clusters, we computed markers for each cluster, and visualized their expression in a heatmap (Figure 1D, Supplemental Table 1). GC I and GC II clusters shared a fraction of their marker genes, but there was minimal overlap between marker genes of other clusters, indicating, that these cell types are transcriptionally distinct despite shared mesodermal origins. Thus, we produced a high-quality single cell RNA-sequencing dataset of 15,227 cell transcriptomes, which we used to generate ovarian cell type gene expression profiles, transcriptional signatures, and function predictions.

### Identification of 9 distinct steps of GC differentiation

Next, we sought to characterize the transcriptional dynamics underlying GC differentiation. GSCs are in close contact with the niche and are marked by a round GC specific organelle called spectrosome (Lin and Spradling 1995). GSCs divide asymmetrically to give rise to a cystoblast (CB), which subsequently undergoes 4 rounds of synchronous divisions with incomplete cytokinesis to produce a 16-cell cyst. Within the cyst, the spectrosome evolves into a fusome that spans all 16 cells (Huynh 2006). The germarium is divided in four regions organized along the anterior-posterior axis - 1, 2a, 2b and 3 (Figure 2B). GC divisions occur in the region 1, the anterior part of the germarium. Initially, the 16-cell cysts are rounded in shape and multiple can reside next to each other in region 2a of the germarium (Spradling 1993). During these initial stages of GC differentiation, ECs send protrusions around GCs and regulate their differentiation (Schulz et al. 2002, Kirilly et al. 2011, Banisch et al. 2017). As 16-cell cysts transition into region 2b, they flatten to a disc shape and become surrounded by follicle cells. Cysts grow larger and rounder as they reach region 3, where stage 1 of oogenesis begins. Finally, a follicle containing a cyst surrounded by epithelial FCs pinches off from the germarium and transitions through 14 stages of oogenesis to give rise to a mature egg (Figure 2B). One of the 16 cells is specified as an oocyte and is destined to become the egg, while the other 15 cells become nurse cells that support oocyte growth by providing mRNAs and proteins. Towards the end of oogenesis during stage 11, nurse cells transfer all their content to the oocyte and subsequently undergo apoptosis (Spradling 1993).

**Figure 2.**
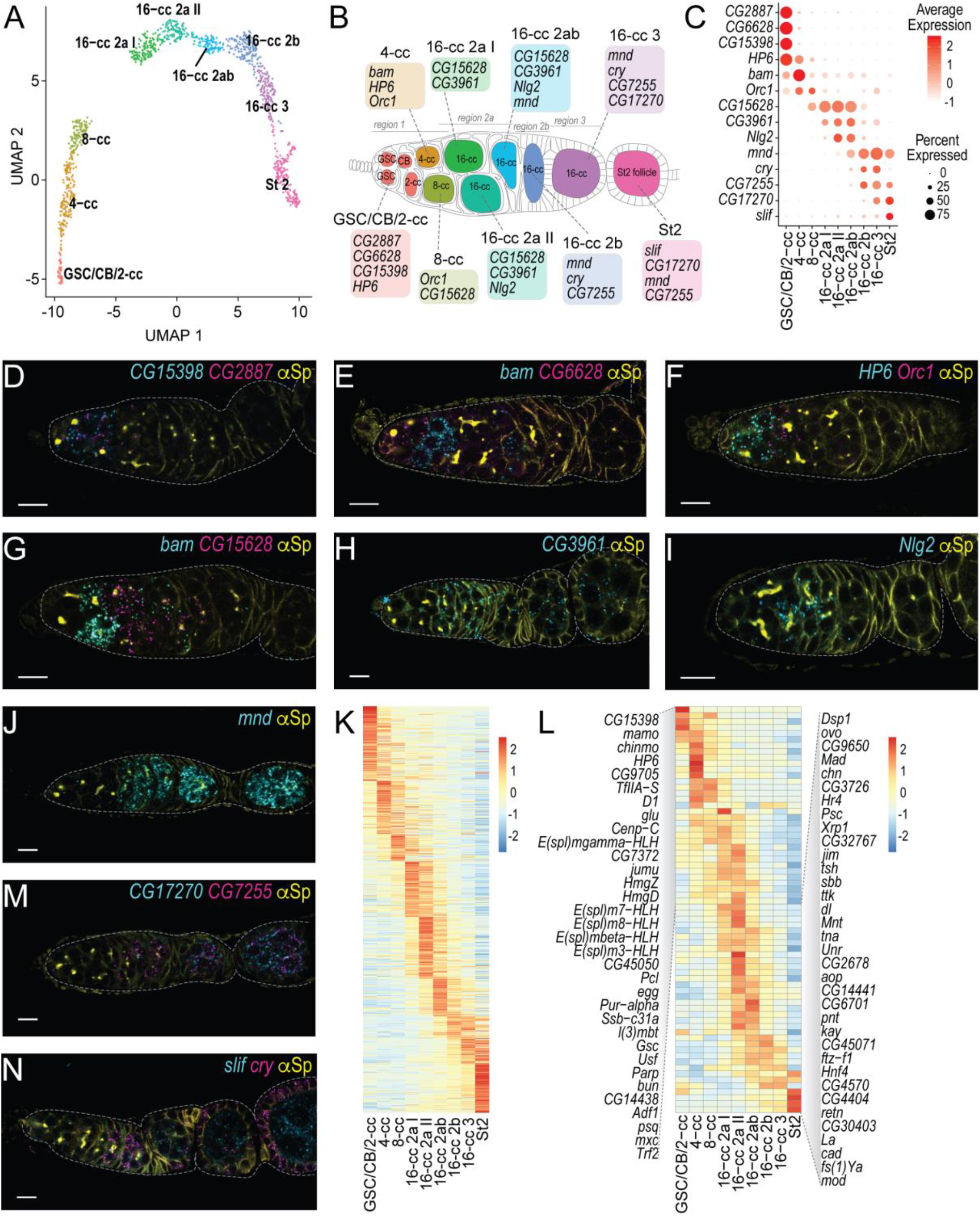
GC subclustering in 9 differentiation states. A - UMAP plot of GCs subclustered in 9 differentiation states. Each dot represents a transcriptome of a single cell, and is color-coded according to cluster membership. B - schematic drawing of a germarium and stage 2 follicle with GCs highlighted in the colors corresponding to the UMAP plot in 2A. Differentiation stage (GSC, CB, and cyst stage) are indicated in each cell. TFs, CCs, ECs and FC lineages are colored in white. Germarium regions are indicated above the cartoon. Color-coded boxes indicate select marker genes for each cluster, whose mRNAs are visualized in 3D-J, 3M, 3N. C - A dot plot visualizing expression of 14 marker genes in 9 GC clusters. Dot diameter represents the fraction of cells expressing each gene in each cluster, as shown in scale. Color intensity represents the average normalized expression level. D-J, M, N - Marker gene mRNA *in situ* hybridization using HCR (cyan, magenta). Spectrosomes, fusomes and somatic cell membranes are labeled by anti-α-Spectrin (yellow). Scale bars −10 μm. D - *CG15398* (cyan) and *CG2887* (magenta) are expressed in GSCs, CBs and 2-cell cysts. E - *bam* (cyan) is predominantly expressed in 4-cell cysts, *CG6628* (magenta) is expressed in GSCs, CBs and 2-cell cysts. F - *HP6* (cyan) is expressed in GSCs, CBs, 2-cell and 4-cell cysts, *Orc1* (magenta) is expressed in 4-cell and 8-cell cysts. G - *bam* (cyan) is predominantly expressed in 4-cell cysts, with lower expression in 2-cell and later stage cysts, *CG15628* (magenta) is expressed in 8-cell and 16-cell cysts in germarium regions 2a and 2b. H - *CG3961* (cyan) is expressed predominantly in 16-cell cysts in germarium regions 2a and 2b. I - *Nlg2* (cyan) is expressed in 16-cell cysts in germarium regions 2a and 2b. J - *mnd* (cyan) is expressed in 16-cell cysts in germarium regions 2a, 2b and 3, and in stage 2 follicle. K, L - A heatmap visualizing average gene expression levels of all GC marker genes (K) and dynamically expressed transcription factors (L) in each cluster. Red indicates highest and blue lowest expression. M - *CG17270* (cyan) is expressed in 16-cell cysts in germarium region 3 and in stage 2 follicle, *CG7225* (magenta) is expressed in 16-cell cysts in germarium regions 2b and 3, in stage 2 follicle and beyond. N - *slif* (cyan) is expressed in stage 2 follicle and beyond, *cry* (magenta) is expressed in 16-cell cysts in germarium regions 2b and 3, and in follicle cells.

**Figure 3.**
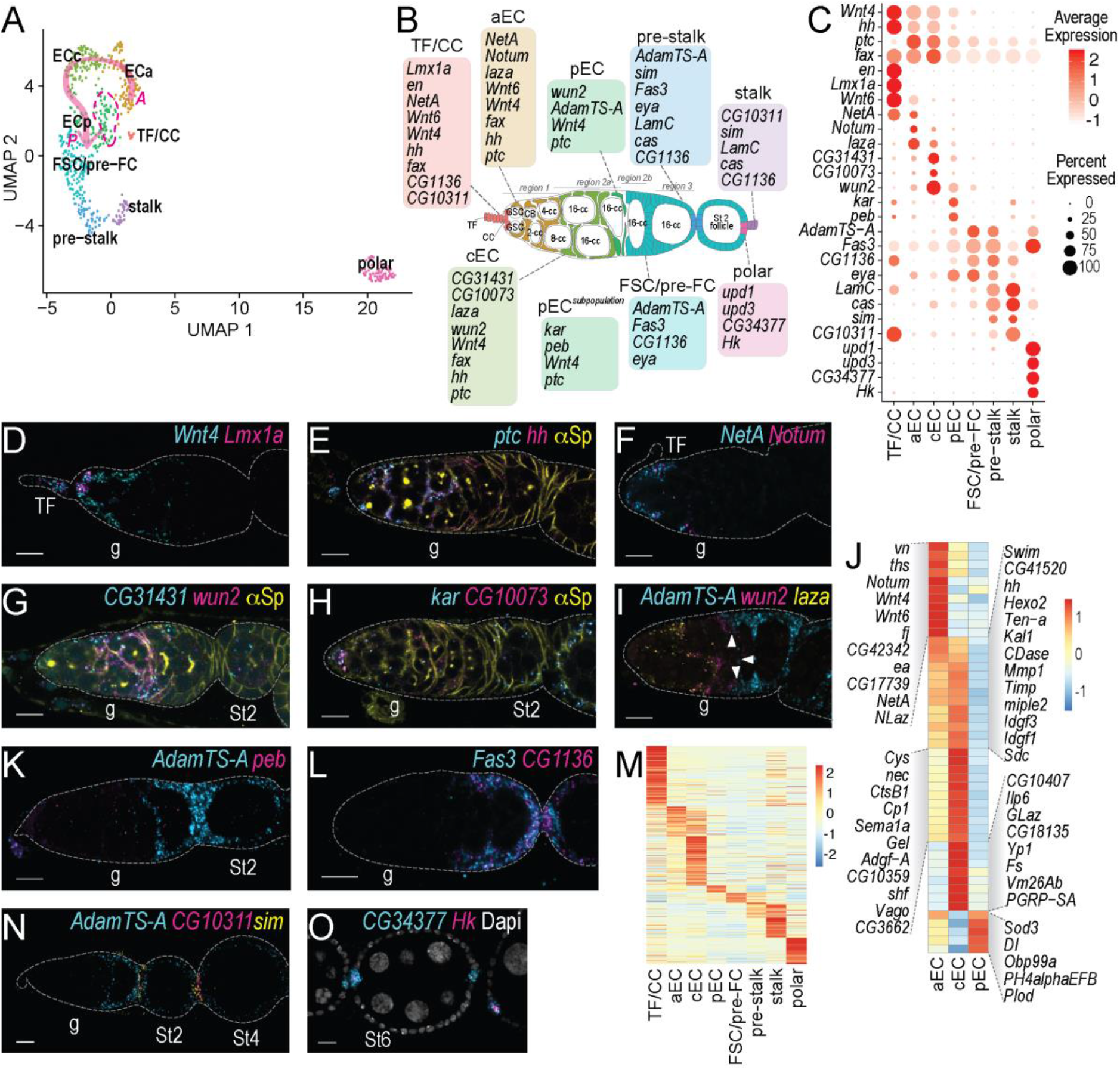
Identification of three transcriptionally distinct EC subtypes. A - UMAP plot of somatic cells of the germarium subclustered in 8 clusters. Each dot represents a transcriptome of a single cell, and is color-coded according to cluster membership. B - schematic drawing of a germarium and stage 2 follicle with somatic cells highlighted in the colors corresponding to the UMAP plot in 3A. GC are colored in white, and their differentiation stage (GSC, CB, and cyst stage) are indicated in each cell. Germarium regions are indicated above the cartoon. Color-coded boxes indicate select marker genes for each cluster. C - A dot plot visualizing expression of select 28 marker genes. Dot diameter represents the fraction of cells expressing each gene in each cluster, as shown in scale. Color intensity represents the average normalized expression level. D-I, K-L, N-O - Marker gene mRNA *in situ* hybridization using HCR (cyan, magenta, and yellow). Scale bars, 10 μm. g - indicates germaria, TF - terminal filament, St2, St4, and St6 indicates oogenesis stages for follicles. E, G, H - Spectrosomes, fusomes and somatic cell membranes are labeled by anti-α-Spectrin (yellow). D - *Wnt4* (cyan) is expressed in TF, CC and ECs, *Lmx1a* (magenta) is expressed in TF and CC. E *ptc* (cyan) is expressed in ECs, *hh* (magenta) is expressed in ECs and pre-FCs. F - *NetA* (cyan) and Notum (magenta) are expressed in anterior ECs in germarium region 1. G - *CG31431* (cyan) is predominantly expressed in EC in germarium region 2a with some expression in region 1 and 2b, *wun2* (magenta) is predominantly expressed in EC in germarium region 2a. H - *kar* (cyan) is sparsely and lowly expressed throughout the germarium, *CG10073* (magenta) is predominantly expressed in region 2a ECs. I - *AdamTS-A* (cyan) is expressed in FSC, pre-FCs, and the most posterior ECs, *wun2* (magenta) is predominantly expressed in EC in germarium region 2a, *laza* (yellow) is expressed in anterior ECs in germarium region 1 and some ECs in germarium region 2a. J - A heatmap visualizing average expression levels of EC enriched secreted proteins and ECM components. Red indicates highest and blue lowest expression. K - *AdamTS-A* (cyan) is expressed in FSC, pre-FCs, and the most posterior ECs, *peb* (magenta) is sparsely and lowly expressed throughout the germarium. L - *Fas3* (cyan) and *CG1136* (magenta) are expressed in FC lineage including FSCs, pre-FCs, pre-stalk, and epithelial FCs. M - A heatmap visualizing average gene expression levels of all somatic cell marker genes in each cluster. Red indicates highest and blue lowest expression. N - *AdamTS-A* (cyan) is expressed in FC lineage including FSCs, pre-FCs, pre-stalk, and epithelial FCs, *CG10311* (magenta) is expressed in stalk cells, *sim* (yellow) expressed in pre-stalk and stalk cells. O - *CG34377* (cyan) and *Hk* (magenta) are expressed in polar cells, Dapi (white) labels cell nuclei.

To identify gene expression profiles specific to distinct stages of GC differentiation, we sub-clustered the GC I and GC II clusters and obtained 9 clusters (Figure 2A). To determine the identity of each cluster, we identified cluster specific marker genes (Figure 2C, Supplemental Table 1), selected markers that are not or are lowly expressed in other cell types (Supplemental Figure S2), and visualized their expression patterns using hybridization chain reaction (HCR), a highly sensitive fluorescence *in situ* hybridization (FISH) method (Choi et al. 2018).

The cluster expressing *CG2887*, *CG6628,* and *CG15398* corresponded to the earliest stages of GC differentiation and contained GSCs, CBs, and 2-cell cysts (cluster GSC/CB/2-cc) (Figures 2B-E). *bam*, a master regulator of GC differentiation onset, was lowly expressed in 2-cell cysts, displayed the strongest expression in 4-cell cysts (cluster 4-cc), and lower levels at later stage cysts (Figure 2B-C, 2E, 2G). *HP6* was expressed in GSCs and early cysts (clusters GSC/CB/2-cc and 4-cc), while *Orc1* was expressed in 4- and 8-cell cysts (clusters 4-cc and 8-cc) and absent from GSCs and 2-cell cysts (Figures 2C, 2F). Thus, *HP6* and *Orc1* co-expression in 4-cell cysts demarcated cluster 4-cc. *CG15628* was expressed starting from 8-cell cyst stage demarcating cluster 8-cc, and continued until 16-cell cysts reached region 2b (Figures 2B, 2C, 2G). The next five clusters corresponded to 16-cell cysts at various stages. Expression of *CG3691* and *Nlg2* was first detected in 16-cell cysts in region 2a, thus demarcating clusters 16-cc 2a I and 16-cc 2a II (Figures 2B, 2C, 2H, 2I). The next cluster contained 16-cell cysts from region 2a and 2b (16-cc 2ab). It was marked by the onset of *mnd* expression, which was first detected in 16-cell cysts in region 2a, and by *CG3961* whose expression extends into 16-cell cysts in region 2b (Figures 2B, 2C, 2H, 2J). Robust expression of *CG7255* was first detected in 16-cell cysts in region 2b marking cluster 16-cc 2b (Figures 2B, 2C, 2M). Finally, *CG17270* expression onset in 16-cell cysts in region 3 marked the cluster 16-cc 3, and *slif* expression onset marked stage 2 follicles (cluster St2) (Figure 2B, 2C, 2M, 2N). *cry* was predominantly expressed in 16-cell cysts in regions 2b and 3 (clusters 16-cc 2b and 16-cc 3) and was downregulated in stage 2 follicles (cluster St 2) (Figures 2B, 2C, 2N). Thus, by exploring the expression of 14 marker genes *in situ*, we have assigned all 9 clusters to specific stages of GC differentiation. The clusters were arranged on the UMAP in the order of their differentiation status, with undifferentiated cells at the bottom left corner, and differentiated cells on the right side (Figure 2A). This indicates that differentiation is a predominant source of transcriptome variation in our dataset. We were unable to clearly distinguish nurse cells from oocytes at the stages examined, likely due to the low number of oocytes in our analyses and the shared cytoplasm between these cell types. GCs from later stages of oogenesis were not present in our dataset due to their larger size, and thus, incompatibility with our cell suspension preparation and cell capture methods.

We were unable to clearly distinguish the GSCs from CBs and 2-cell cysts by clustering. This is likely due to the high degree of similarity in the gene expression profiles of GSCs, CBs, and 2-cell cysts, and the reliance on translational rather than transcriptional control for early GSC differentiation (Slaidina and Lehmann 2014), and due to low abundance of these cell types. An observational study using wild-type ovaries estimated that in each germarium, on average there were 2 GSCs, 1.3 CBs, 1.6 2-cell, 0.9 4-cell, 0.8 8-cell cysts, and 8.4 16-cell cysts. Of the 16-cell cysts, 5.5 were found in region 2a, 1.9 in region 2b, and 1 in region 3 (Drummond-Barbosa and Spradling 2001). Our analysis revealed distinct transcriptomes for 9 steps of GC differentiation from GSC to stage 2 of oogenesis and identified distinguishing markers for most (Figure 2B), suggesting that the transcriptomes identified may correspond to the developmental stages described previously.

### Gene expression dynamics during GC differentiation

Next, we characterized gene expression dynamics during GC differentiation more closely. 813 genes were differentially expressed between GC clusters, indicating that even though the first steps of differentiation rely heavily on translational regulation, the subsequent steps involve transcriptional regulation (Slaidina and Lehmann 2014). We visualized the mean expression levels of differentially expressed genes on a heatmap (Figure 2K, Supplemental table 1, Supplemental table 2). 16-cell cyst clusters displayed remarkably distinct transcriptional signatures. The transcriptional differences between the five germarial 16-cell cyst states indicates that 16-cell cysts become transcriptionally diverse as they are transiting from the proliferation to differentiation. In region 2a 16-cell cysts are enwrapped by ECs, while in region 2b and 3, GCs are surrounded by pre-FCs, which start forming the follicular epithelium. Thus, transcriptional differences may also reflect the changing signals GC cysts receive from various bordering somatic cell types.

Next, we explored the functional predictions for these genes using Gene List Annotation for *Drosophila* (GLAD) (Hu et al. 2015) (Figure S1D, Supplemental Table 3). All GCs in contact with ECs (up to 16-cell cysts in region 2a) were enriched for transmembrane proteins, some of which may be involved in signaling between GCs and ECs, and others in formation of the fusome, which is composed of ER derived vesicles (Huynh 2006). Furthermore, the GSCs/CB/2-cc cluster was enriched for chaperones and heat shock proteins, which have a protective function. Consistent with divisions of early cysts, GSC/CB/2-cc and 4-cc clusters were enriched for cytoskeletal proteins, in particular, those involved in cytokinesis. In contrast, most late stage cysts were enriched for mitochondria related genes, which coincide with rapid expansion of the mitochondrial pool, and selection based on their fitness (Hill et al. 2014, Lieber et al. 2019). Finally, transcription factors and DNA binding proteins were significantly enriched at multiple stages over the course of GC differentiation, and we visualized their expression in a heatmap (Figure 2L). A few were expressed in GSC/CB/2-cc cluster, including, *CG15398* (Figure 1D) and *chinmo* which regulates cyst stem cell renewal in *Drosophila* testis (Flaherty et al. 2010), suggesting that it may have conserved role in stem cell renewal across many tissues. Five Notch responsive *Enhancer of split Complex* (*E(spl)-C*) transcription factors were expressed in 4-cc, 8-cc and 16-cc 2a I clusters, suggesting that Notch signaling pathway may regulate early steps in GC differentiation.

Altogether, we have identified specific transcriptional signatures for 9 distinct steps of GC differentiation. Further analyses of gene expression dynamics between the clusters may reveal novel aspects of GC differentiation regulation.

### Drosophila germaria house three EC subtypes

Next, we aimed to identify transcriptional signatures for somatic cells of the germarium. There are five major somatic cell types in germaria. At the most anterior tip of the germarium reside the somatic terminal filament (TF) and cap cells (CC) of the GSC niche. Directly posterior to them in regions 1 and 2a are the ECs, which extend protrusions that enwrap the GCs. Posterior to the ECs at the region 2a and 2b boundary are the FSCs, which divide asymmetrically to give rise to a transit amplifying FC population called pre-follicle cells (pre-FC). These pre-FCs reside in regions 2b and 3 where they proliferate and further differentiate into polar, stalk and epithelial FC lineages.

To assign transcriptomes to individual cell types, we subclustered the germarium somatic cell cluster and initially obtained seven clusters. We noticed, however, that a small number of cells located separately on the UMAP were included in one of the larger clusters by the clustering algorithm. These cells expressed *en* and *Lmx1a,* well known TF and CC markers (Bolívar et al. 2006, Allbee et al. 2018) (Figure 3A-D, S1E). Therefore, we manually annotated this cell population as TF/CCs. Each ovariole contains only about 8 TFs and 6 CC. In total, our dataset contained only 10 such cells, and thus we were unable to distinguish TFs from CCs. We computed markers for all 8 clusters (Supplemental Table 1). Of the remaining seven clusters, three expressed high levels of the EC markers *fax* and *Wnt4* (Decotto and Spradling 2005, Mottier-Pavie et al. 2016), suggesting that they correspond to three subtypes of ECs (Figure 3A-D), while the other four correspond to FC subtypes expressing *Fas3*, *cas*, or both (Figure 3L).

*hh*, *Wnt4* and *ptc* are expressed in an anterior to posterior gradient, and their expression differed between the EC clusters; we thus hypothesized that the three clusters might correspond to distinct anatomical positions – anterior (aEC), central (cEC), and posterior (pEC) (Figure 3A-E). Moreover, *Wnt6* functions in anterior ECs (Wang and Page-McCaw 2018), and was enriched in the aEC compared to other EC clusters (Figure 3C). We selected a few EC subtype markers, and assessed their mRNA expression in the ovary. *NetA* and *Notum* were specifically expressed in the aEC cluster (Figure 3C), and indeed labeled anterior ECs in region 1 (Figure 3F). *CG31431* and *CG10073* were expressed in the cEC cluster (Figure 3C), and their mRNAs were detected in the central region of the germarium, in close contact to 8- and 16-cell cysts (Figure 3G, 3H). We used a combination of two markers to identify the remaining EC population. *wun2* is expressed in cEC and pEC clusters, while *AdamTS-A* is expressed in the pEC and FC lineage (Figure 3C); therefore, cells co-expressing *wun2* and *AdamTS-A* correspond to pEC. We observed that *wun2* is expressed in ECs in Region 2a, and *AdamTS-A* is the early FC lineage, and expression of both genes overlapped near region 2a and 2b boundary (Figure 3I, arrowheads). These results suggest that the aEC, cEC, and pEC clusters correspond to the most anterior, central and posterior ECs. EC transcriptomes are aligned from anterior to posterior as indicated by a pink arrow in the UMAP plot (Figure 3A, pink arrow).

Two additional pEC markers, *kar* and *peb*, were expressed in a fraction of pEC that lacked expression of *wun2* and *AdamTS-A* (Figure 3A, dotted line in magenta, 3C, Supplemental Figure S1E). In the germarium, *kar* and *peb* displayed sparse weak expression throughout the germarium (Figure 3H, 3K), suggesting that there might be additional complexity within EC population that has not been revealed by clustering. To test whether we captured all EC subtypes, we labeled *laza*, which is expressed in aEC and cEC together with *wun2* and *AdamTS-A*, and observed that all regions of the germarium where labeled, except the most anterior tip (TF/CC) (Figure 3I). Thus, we have, indeed, captured all EC subtypes.

Altogether, we have identified three transcriptionally distinct EC subtypes arranged along the anterior-to-posterior axis of the germarium. To explore the potential functional differences between the EC subtypes in our dataset, we compared their transcriptomes. 405 genes were differentially expressed between EC clusters. We visualized their expression in a heatmap together with other somatic cell markers (Figure 3M) and performed GLAD analyses on EC marker genes (Figure S1F, Supplemental Table 3). Consistent with their role as GC differentiation regulators, ECs were strongly enriched for gene classes related to cell-cell communication, like, signaling pathway components, secreted proteins, receptors, transmembrane proteins, and extracellular matrix (matrisome) proteins.

Because of their anatomical position, the three EC populations are in contact with GCs at different states of differentiation, suggesting that each EC subtype might send a distinct set of signals to regulate different stages of GC self-renewal or differentiation. To identify such potential regulators, we visualized the expression of all secreted proteins and ECM components that are differentially expressed among ECs subtypes (Figure 3J). *Fs* (*Follistatin*), an inhibitor of activin and *dpp* signaling (Bickel et al. 2008), was expressed specifically in cECs. Since *dpp* signaling promotes GSC self-renewal and impedes GC differentiation, it is plausible that *Fs* expression in the cECs may facilitate GC differentiation by suppressing the *dpp* signal emanating from the GSC niche.

aECs expressed Wnt signaling pathway components and regulators, including ligands *Wnt4* and *Wnt6*, and *Notum,* which regulates Wnt ligand activity (Kakugawa et al. 2015). aECs expressed also EGF and FGF ligands *vn* and *ths.* Insulin-like peptide *Ilp6* was specifically expressed in cEC, and two regulators of insulin/IGF signaling *Idgf1* and *Idgf3* were expressed in aEC and cEC. Thus, a number of signaling pathway activators and regulators were expressed in distinct regions of the germarium and may regulate GC differentiation. Alternatively, ECs may signal to other somatic cells of the germarium, like, TFs and CCs, or FSCs. Further studies will discriminate between these possibilities. cECs expressed also a number of secreted peptidases (*Cys*, *CtsB1*, *Cp1*), which may be involved in signaling molecule activation or degradation. Moreover, peptidase inhibitors *nec*, *CG17739* and *Kal1* were expressed in adjacent EC clusters and, thus may refine the peptidase activity domain. pECs were enriched for genes required for collagen production and secretion (*PH4alphaEFB*, *Plod*), while aEC and cEC expressed matrix metalloprotease (*Mmp1*) and its inhibitor (*Timp*), which regulate ECM structure, suggesting that there is distinct ECM structure and composition in different parts of the germarium. Moreover, a number of adhesion molecules and axon guidance molecules (*NetA*, *Ten-a*, *Sema1a*) where differentially expressed between the EC subtypes. Altogether, these results indicate that each EC subgroup secretes a different set of molecules, which may create distinct microenvironments for GC differentiation along the anterior-posterior axis of the germarium. Since signaling between ECs and GCs relies at least in part on large surface contacts between GCs and EC protrusions (Schulz et al. 2002), we sought to search for physical interactor pairs (such as transmembrane-transmembrane and secreted-transmembrane) that may mediate signaling between GCs and ECs (Guruharsha et al. 2011). We selected secreted ligands, extracellular matrix proteins (ECM), and transmembrane proteins enriched in EC (158 genes) or GC (200 genes) clusters adjacent to ECs (GSC/CB/2-cc, 4-cc, 8-cc, 16-cc 2a I, 16-cc 2a II, and 16-cc 2ab). 49 genes (16 expressed in GCs, 23 in ECs, and 10 in both) formed 119 predicted protein: protein interaction pairs between transmembrane/secreted proteins in GCs and ECs (Supplemental Table 4). In 30 pairs, both genes had GO terms associated to plasma membrane or extracellular space, and in 65 pairs at least one of the genes was uncharacterized (in 17 both were uncharacterized). Further analyses of the physical interaction pairs may reveal novel mechanisms for GC differentiation regulation by ECs.

Altogether, our data together with a recent report by Rust *et al*. identifies three EC subtypes and suggests functional specialization among ECs. Subtype specific markers, transcriptomes, and potential signaling molecules and interactors provide a resource for the functional characterization of EC subtypes and may reveal novel signaling mechanisms between GCs and soma.

### Identification of early FC lineages

We next sought to characterize the FC lineages present in the germarium soma cluster. One cluster was rather distant from other clusters and expressed polar cell markers *upd1* and *upd3* (Figure 3A-C). Visualization of two newly identified markers *CG34377* and *Hk*, confirmed that the cluster corresponded to polar cells (cluster polar) (Figure 3C, 3O).

Stalk cells express *LamC* and *cas* (Song and Xie 2003, Chang et al. 2013). Two adjacent clusters expressed both markers, albeit at different levels (Figure 3A, 3C). To determine the cluster identity, we searched for genes that were enriched and differentially expressed between the clusters. *sim* was expressed in both, while *CG10311* and *AdamTS-A* were each enriched in one of the clusters (Figure 3C). *sim* was expressed in FCs at region 3 and the stage 2 follicle interface, and in stalk cells (Figure 3N), suggesting that the two clusters correspond to stalk cells and FCs differentiating into stalk cells (pre-stalk). *CG10311* was strongly expressed in stalk and weakly expressed in pre-stalk, while *AdamTS-A* was expressed in pre-stalk (and other pre-FCs), but absent from stalk cells, indicating that the *AdamTS-A* expressing cluster corresponds to the pre-stalk cells, and *CG10311* expressing cluster corresponds to the stalk cells (Figure 3C, 3N).

Finally, the remaining cluster expressed high levels of *AdamTS-A* and *CG1136*, which are both expressed in pre-FCs in germarium regions 2b and 3 (Figure 3K, 3L). Therefore, we concluded that this cluster contains FSCs and pre-FCs (FSC/pre-FC). Consistently, this cluster expressed also high levels pre-FC marker *eya* (Figure 3C) (Bai and Montell 2002). Our clustering analyses did not reveal a clear FSC population, likely because the low numbers of FSCs per germarium resulted in isolation of too few FSC transcriptomes for our analyses (Fadiga and Nystul 2019, Reilein et al. 2017).

Aiming to identity the FSCs and their markers, we performed pseudotime analyses on the follicle cell lineages (clusters FSC/pre-FC, pre-stalk, stalk, polar) using Monocle, an excellent tool to align cells on a “pseudotime” axis and study gene expression dynamics over the course of differentiation (Qiu et al. 2017). These analyses revealed two branches of FC differentiation, one corresponding to polar cells and the other to stalk cells, where FCs transition through a pre-stalk state before acquiring stalk cell identity (Figure S3A-B). *eya* was downregulated as FCs committed to polar and stalk cell fate (Figure S3C). In contrast, *upd1* was upregulated in polar cells. *CG10311* was upregulated and *AdamTS-A* was downregulated in stalk cells over time (Figure S3D-F). Thus, our analyses are in agreement with the literature (Tworoger et al. 1999, Beccari et al. 2002) and with our cluster assignment to cell types, and we therefore used it for identification of putative FSCs and their markers.

We identified two putative markers for FSCs, *if* and *CG6044* (Figures S3G-I), and visualized their expression using HCR. In addition, we labeled *Fas3* mRNA, which labels follicle cell lineage, and Dapi, which labels cell nuclei, to discern the precise location of region 2a/2b boundary where the FSCs are positioned. We found that *if* was expressed in the region where FSCs reside, however its expression extended to pre-FCs and ECs, where it was weakly expressed (Figure S2J). *CG6044* was predominantly expressed in the pre-FCs, with weak expression at the germarium region 2a and 2b boundary where FSCs reside (Figure S2K). Thus, we were unable to identify FSC specific markers, likely because the FSC gene expression profile is highly similar to the pre-FC gene expression profiles (Nystul and Spradling 2010, Hartman et al. 2015), and therefore they are indistinguishable bioinformatically in our dataset. Increasing the number of cells of interest might increase the statistical power enough to alleviate these limitations.

Rust *et al.* used a similar approach to identify FSCs and their markers. While their markers are indeed enriched in our putative FSC population, their expression extends to ECs and pre-FCs similarly to the markers we identified (Supplemental Text).

399 genes were differentially expressed between the FC clusters, and we visualized their expression in a heatmap (Figure 3M). The polar cell cluster displayed the most distinct signatures, while pre-stalk and stalk cells shared a fraction of their marker genes. We performed GLAD analyses on all marker genes (Figure S1F). Major signaling pathway components were enriched in FC lineages, consistent with high signaling activity during FC lineage specification. FC lineage clusters were also enriched for cytoskeletal genes. The FSC/pre-FSC cluster was particularly enriched for cell division related cytoskeletal proteins, consistent with active proliferation. A number of TF/CC specific genes were also enriched in stalk cells. Both cell types are post-mitotic and share a similar flattened shape, suggesting that many of the shared genes may comprise cell shape regulators. Altogether, we determined FC lineage gene expression profiles and markers. Exploration of the gene expression dynamics as FSCs transition into specialized FC fates may reveal novel regulatory mechanisms for specification of FC lineages.

### Identification of epithelial FC subtypes

Epithelial follicle cells together with GCs transition through 14 stages of oogenesis and are the most abundant ovarian cell type. FC and GC progression through oogenesis is highly coordinated, since FCs play an essential role in egg development by determining the body axes of the future embryo and producing essential yolk proteins (Wu et al. 2008). Determining the transcriptional differences between FC subtypes will strengthen our understanding of how this coordination is achieved. Because we manually removed late stage follicles during dissections (see Materials and Methods), we expected to identify FC transcriptomes from early and mid-stage follicles. Indeed, stage 9/10A FC marker *Yp2* was well expressed, but stage 10B marker *Cp36* expression was barely detectable in our dataset (Supplemental Figure S4A) (Tootle et al. 2011), indicating that our dataset contains FC transcriptomes up to stage 9/10A. Therefore, we focused on FC development up to that stage.

Follicle cells progress from mitotic (stage 1-6) to transitioning (stage 7) to endocycling states (stage 7 - 9) (Deng et al. 2001) (Figure 4D). Starting from stage 5, FCs differentiate into multiple subtypes determined by external cues, which largely depend on the anatomical position within the follicle (Wu et al. 2008). FCs at the anterior and posterior tip of the follicle receive JAK/STAT signal (*upd1* and *upd3*) from polar cells, and assume terminal fate, while the rest become mainbody (MB) follicle cells (MBFC) (Xi et al. 2003). Anterior terminal (AT) follicle cells (ATFC) assume three distinct fates determined by the distance from the tip-border cells, stretched cells, and centripetal cells (Xi et al. 2003). Posterior terminal cells receive an additional EGF signal (*grk*) emanating from the oocyte nucleus starting from stage 6 and assume the posterior terminal (PT) follicle cell (PTFC) fate (González-Reyes et al. 1995). Initially, epithelial FCs have cuboidal shape, and, at stage 9, the PTFCs transition into columnar shape FCs. Simultaneously, border cells together with anterior polar cells start migrating posteriorly between the nurse cells. Stretched cells start producing a TGF-beta ligand (*dpp*), and in response to it flatten in a wave from the anterior to posterior covering the rapidly growing nurse cells (Brigaud et al. 2015). The MBFCs cover the growing oocyte and assume columnar shape as well. At stage 9/10A, oocyte nucleus migrate to the anterior dorsal corner of the oocyte, and by activating EGFR signaling specifies the dorsal FCs (Spradling 1993, González-Reyes et al. 1995). Centripetal cells, which initially are located between stretched cells and MBFCs, start migrating between the nurse cells and oocyte at stage 10B (Wu et al. 2008). Further follicle cell morphogenesis events occur beyond this stage, and each follicle cell subtype has a specific role in forming various structures of eggshell (Duhart et al. 2017).

**Figure 4.**
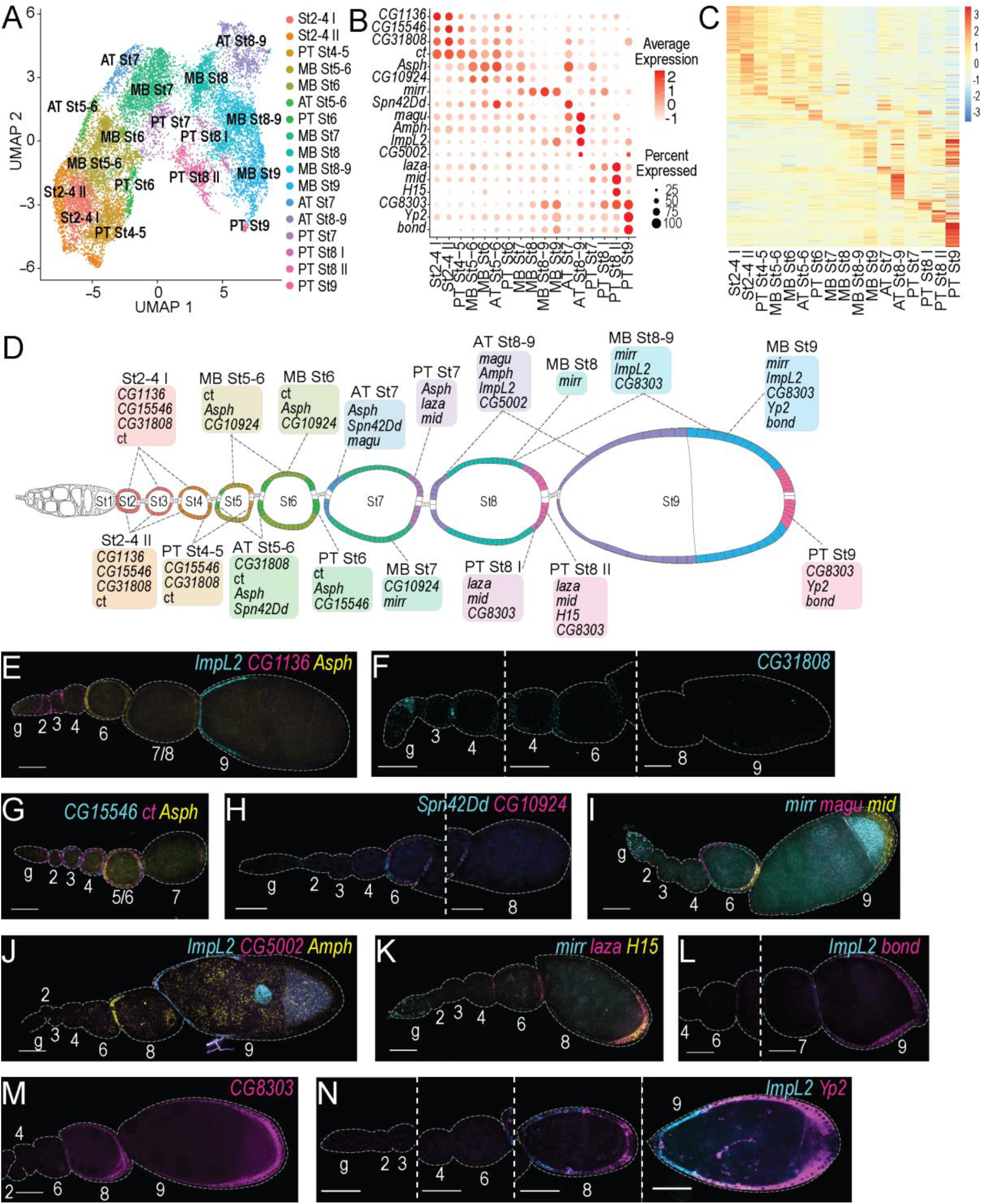
Characterization of 17 FC subpopulations. A - UMAP plot of FCs subclustered in 17 clusters. Each dot represents a transcriptome of a single cell, and is color-coded according to cluster membership. B - A dot plot visualizing expression of select 18 marker genes. Dot diameter represents the fraction of cells expressing each gene in each cluster, as shown in scale. Color intensity represents the average normalized expression level. C - A heatmap visualizing average gene expression levels of all FC marker genes in each cluster. Red indicates highest and blue lowest expression. D - schematic drawing of an ovariole with FC subtypes highlighted in the colors corresponding to the UMAP plot in 4A. E-N - Marker gene mRNA *in situ* hybridization using HCR (cyan, magenta, yellow). Follicle stages (2-9) and germarium (g) are indicated above or below each follicle. Scale bars, 30 μm. E - *ImpL2* (cyan) is expressed in stretch cells at stage 9, *CG1136* (magenta) is expressed in pre-FCs, and stage 2-4 FCs, *Asph* (yellow) expressed in stage 6 FC, and restricted to both termini at stage 7. F - *CG31808* (cyan) is faintly expressed in stage 3-4 FCs, and is restricted to ATFC at stage 6. G - *CG15546* (cyan) has patchy expression in stage 2-4 follicles, and is restricted to PTFC by stage 5-6, *ct* (magenta) is expressed in all FCs from pre-FC to stage 6, *Asph* (yellow) expressed in stage 5-6 follicle. H - *Spn42Dd* (cyan) is expressed in ATFCs at stage 6, *CG10924* (magenta) is expressed in MBFCs at stage 6. I - *mirr* (cyan) is expressed in MBFC from stage 6-7 onwards, *magu* (magenta) is expressed in ATFCs from stage 6 and is downregulated by stage 9, *mid* (yellow) expressed in PTFC starting from stage 6. J - *ImpL2* (cyan) is expressed in ATFC excluding the most anterior tip at stage 8 and in stretch and border cells at stage 9, *CG5002* (magenta) is expressed in stretch cells at stage 9, *Amph* (yellow) expressed in ATFC at sage 8. K - *mirr* (cyan) is expressed in MBFC from stage 6-7 onwards, *laza* (magenta) is weakly expressed in PTFCs starting from stage 4, and strongly expressed in a broad band of PTFC at stage 8, *H15* (yellow) is expressed in a narrow band of the most posterior PTFCs at stage 8, weak expression is detectable in PTFCs at stage 6. L - *ImpL2* (cyan) is expressed in stretch cells at stage 9, *bond* (magenta) is expressed in a posterior-to-anterior gradient in PTFCs and MBFCs at stage 9. M - *CG8303* (magenta) is expressed in a posterior-to-anterior gradient in PTFCs and MBFCs at stages 8 and 9. N - *ImpL2* (cyan) is expressed in ATFCs at stage 8, extending into MBFCs, and in stretched cells at stage 9, *Yp2* (magenta) is expressed in a posterior-to-anterior gradient in PTFCs and MBFCs at stages 8 and 9. Its expression is excluded from the most posterior tip.

To find gene expression profiles of epithelial follicle cell subpopulations, we performed in-depth analyses on FC I, II and III clusters. We first assessed expression of known markers to assign transcriptomes to distinct stages of oogenesis and follicle cell subtypes.

For staging, we assessed the expression of *ct*, *stg*, *CycB* and *peb*. *ct* and *CycB* are expressed in mitotic FCs up to stage 6, while *stg* expression extends into stage 7 (Figure 4D, 4G, Supplemental Figure S4A) (Jackson and Blochlinger 1997, Sun and Deng 2005, Deng et al. 2001). Transcription factor *peb* (also called *Hindsight*) is expressed sporadically starting at stage 6 and its expression is strongly upregulated during stage 7 and beyond (Supplemental Figure S4A) (Sun and Deng 2007). As mentioned above, *Yp2* marks stage 9 follicles (Figure 4D, 4N, Supplemental Figure S4A) (Tootle et al. 2011). Therefore, the end of stage 6 is marked by downregulation of *CycB* and *ct*, stage 7 is marked by overlapping expression of *stg* and *peb* (Supplemental Figure S4A), *peb* expression in absence of *stg* marks FCs starting from stage 8, and *Yp2* marks FCs starting from stage 9. Therefore, in the UMAP plot (Figure 4A), transcriptomes of the early stage FCs are located in the bottom left corner and transition in an arch towards the right where FCs differentiate into multiple subtypes.

Next, we sought to assess whether the anatomical position correlated with specific UMAP regions. We assessed expression patterns of a number of MB and terminal follicle cell markers. *mirr* marks MBFCs, and was expressed in a band in the middle of the UMAP (Figure 4A, 4I, 4K, Supplemental Figure S4A) (Jordan et al. 2000). *mid*, *H15* and *pnt* are expressed at posterior terminus at stage 6 and onwards, and we observed their expression in a band on the bottom of UMAP (Figure 4A, 4I, 4K Supplemental Figure S4A) (Morimoto et al. 1996, Lomas et al. 2013). Following dorsal migration of oocyte nucleus at stage 9, *pnt*, but not *mid* or *H15,* expression is induced in the anterior dorsal cells (Morimoto et al. 1996, Lomas et al. 2016). In our dataset, *pnt* was always co-expressed with *mid* and *H15*, indicating that the anterior dorsal stage 9/10A cells are absent in our dataset. *dpp* is expressed in stretched cells, and we observed its expression in the top right corner of the UMAP (Supplemental Figure S4A) (Xi et al. 2003). Border cell and centripetal cell marker *slbo* was detected in a very small number of transcriptomes adjacent to *dpp* expressing transcriptomes (Supplemental Figure S4A) (Murphy et al. 1995). Therefore, AT follicle cells are on the top, MB in the middle and PT on the bottom of the UMAP plot (Figure 4A), indicating that the transcriptome variation in FCs is driven by both- the anatomic position in the follicle and the oogenesis stage.

To obtain precise subdivision of FCs by stage and subtype, we subclustered the transcriptomes into 15 clusters, and manually split 2 clusters (Supplemental Text) (Figure 4A). We identified specific markers for each cluster, and visualized their expression along with a few previously described markers (Figure 4B, 4E-N).

*CG1136* was expressed in FCs of very early follicles up to stage 4 [clusters St2-4 I, St2-4 II] (Figure 4B, 4D, 4E, Supplemental Figure S4A). *CG15546* expression was patchy in stage 2-4 follicles [clusters St2-4 I, St2-4 II, PT St4-5], and slightly enriched at the termini prior to enriching to the PT at stage 6 [cluster PT St6] (Figure 4B, 4D, 4G, Supplemental Figure S4A), while *CG31808* had weak patchy expression in stage 2-4 follicles and restricted to AT at stage 5-6 [clusters St2-4 I, St2-4 II, PT St4-5, and AT St5-6] (Figure 4B, 4D 4F, Supplemental Figure S4A). *ct* was expressed in FCs up to stage 6 as previously reported [clusters St2-4 I, St2-4 II, PT St4-5, MB St5-6, MB St6, AT St5-6, PT St6] (Figure 4B, 4D, 4G, Supplemental Figure S4A) (Jackson and Blochlinger 1997). *Asph* was ubiquitously expressed starting from stage 5 and enriched to both termini by stage 7 [clusters MB St5-6, MB St6, AT St5-6, PT St6, AT St7, PT St7] (Figure 4B, 4D, 4E, 4G, Supplemental Figure S4A). MBFCs were labeled by *CG10924* at stage 6-7 [clusters MB 5-6, MB St6, MB St7], and the previously described transcription factor *mirr* from stage 7 onwards [clusters MB St7, MB St8, MB St8-9, MB St9] (Figure 4B, 4D, 4H, 4I, 4K, Supplemental Figure S4A) (Jordan et al. 2000).

We visualized expression of four genes expressed in AT. *Spn42Dd* was predominantly expressed in the AT at stages 5-7 [clusters AT St5-6, AT St7] (Figure 4B, 4D, 4H, Supplemental Figure S4A). *magu* was expressed in AT starting stage 6, peaking at stage 8, and declining at stage 9 [clusters AT St7, AT St8-9] (Figures 4B, 4D, 4I, Supplemental Figure S4A). *Amph* was expressed in AT at stage 8 [cluster AT St8-9], and its expression was undetectable in stretched cells at stage 9 (Figure 4B, 4D, 4J, Supplemental Figure S4A), while *CG5002* was expressed in a very narrow band in stage 8 follicles, and in stretched cells at stage 9 [cluster AT St8-9] (Figures 4B, 4D, 4J, Supplemental Figure S4A). *ImpL2* expression started at stage 8 and was excluded from the very anterior tip but reached far posterior in MBFCs at late stage 8 [clusters AT St8-9, MB St8-9, MB St9]. In stage 9, *ImpL2* was expressed in flattened stretched cells and border cells (Figure 4B, 4D, 4E, 4J, 4L, 4N, Supplemental Figure S4A).

Posterior terminal (PT) FCs were faintly labeled by *laza* and *mid* at early stages, reaching maximum at stage 8 [clusters PT St7, PT St8 I, PT St8 II] (Figures 4B, 4D, 4I, 4K, Supplemental Figure S4A). *CG8303* was enriched in PT at stages 8 and 9 [clusters PT St8 I, PT St8 II, PT St9], *Yp2* and *bond* at stage 9 [cluster PT St 9] but all three were expressed in MBFCs as well [clusters MB St9, and MB St8-9 for CG8303 only] (Figure 4B, 4D, 4L, 4M, 4N Supplemental Figure S4A). Stage 8 PT were divided in two sub-populations: cluster PT St8 II corresponded to the most posterior tip and expressed *H15* along with *laza* and *mid*, and cluster PT St8 I corresponded to a band of cells adjacent to PT St8 II cells, which expressed *laza* and *mid* but not *H15* (Figures 4B, 4D, 4I, 4K, Supplemental Figure S4A).

### Characterization of FC transcriptional signatures

518 genes were differentially expressed between FC subtypes, and we visualized their expression in a heatmap (Figure 4C). We observed temporal signatures (stage 2-6, and stage 8-9) and cell fate dependent (AT, MB, PT) signatures. The gene expression signature of each cluster was composed of blending of both the temporal and cell fate signatures, especially during the earlier stages of oogenesis.

To gain insights of functional differences between the FC subtypes, we explored gene class enrichment in the signature of each cluster (Supplemental Figure S4B-D, Supplemental Table 3) (Hu et al. 2015). Transcription factors/DNA binding proteins were enriched in multiple clusters, highlighting the role of transcriptional regulation in follicle cell subtype diversification. Cytoskeletal proteins were enriched in multiple early stage follicle cell subtypes and in AT at later stages. The sets of genes, however, were distinct. The early proliferative FCs, were enriched for cell division associated cytoskeletal proteins, while stage 7 and 8-9 ATs were predominantly enriched for actin binding proteins and actin regulators, many of which are involved in cell morphogenesis events like axon targeting. In these cells, they are likely involved in stretched cell flattening.

Transporters, in particular, vacuolar ATPase (V-ATPase) subunits were enriched in late stage follicle cells, especially in cluster AT St8-9. V-ATPase is a large multimolecular complex that uses the energy of ATP hydrolyses to create proton gradients, for example, lower the pH of particular cell compartments, like, lysosomes or endosomes (Nelson et al. 2000). V-ATPases locate to stretch cell plasma membrane at stage 13 of oogenesis to acidify nurse cells and facilitate their death and engulfment by stretch cells. It is plausible that FCs begin upregulating V-ATPase subunit genes as early as at stage 8-9 in preparation for the acidification. Indeed, the first signs of nurse cell death can be observed already at stage 10 (Cooley et al. 1992), suggesting that V-ATPases may start executing their function well prior to stage 13.

Numerous additional gene classes were enriched in individual clusters, and future studies may reveal novel functions of specific follicle cell subtypes. Moreover, in-depth studies of differentially expressed transcription factors will help understand the transcriptional networks governing complex FC subtype specification and morphogenesis.

## Discussion

We generated a single cell atlas of the stem cell compartment and early differentiating egg chambers of adult ovaries of *Drosophila melanogaster*. We characterized cell type-specific transcriptional signatures, identified novel markers and generated functional predictions for 34 cell types and subtypes - 9 states of GC differentiation, GSC niche cells, 3 EC subtypes, FSC/pre-FCs, 3 clusters corresponding to polar and stalk cell lineages, and 17 epithelial FC subtypes. This extensive annotation will bolster future studies, for example, by capitalizing on cluster specific markers to develop cell type specific genetic tools.

We could not distinguish GSCs and FSCs from their daughters by clustering. This is likely due to the relatively low numbers of stem cells in our dataset, but it also reflects the high similarity between the transcriptomes of stem cells and their daughters (Kai et al. 2005, Slaidina and Lehmann 2014, Nystul and Spradling 2010, Hartman et al. 2015). This stability of the stem cell transcriptome might have a functional relevance. Indeed, CBs and germline cysts can de-differentiate and compete with GSCs for niche occupancy (Xie and Spradling 1998, Liu et al. 2015). Likewise, FSC daughters migrate across the germarium and compete for niche occupancy with other FSCs (Nystul and Spradling 2007). Thus, stem cell daughters initially retain the ability to revert to a stem cell state, possibly because they do not extensively remodel their transcriptome shortly after the asymmetric division.

ECs have a dual role, they promote GSC self-renewal and GC differentiation forming a domain termed differentiation niche (Kirilly et al. 2011, Wang and Page-McCaw 2018). We and Rust *et al*. identified three EC subtypes - anterior, central, and posterior ECs. Each EC subtype interacts with GCs of a particular differentiation state, and likely sends and receives distinct signals. Likewise, EC morphologies differ between locations; cEC and pEC protrusions are longer than aEC protrusions, as they interact with increasingly large germline cysts (Banisch et al. 2017). We uncovered that a number of secreted proteins, adhesion molecules and ECM components are differentially expressed between EC subtypes, suggesting that each subtype creates a distinct microenvironment. Thus, as GCs progress through differentiation and move posteriorly, their immediate microenvironment changes. These observations open the possibility that the spatial organization of distinct EC microenvironments supports progressive GC differentiation, and that maturing GC may feed back on their microenvironment to define and stabilize its pattern.

Highly granular clustering of GC transcriptomes and precise cluster identity assignments allowed us to identify transcription factors that are dynamically expressed over the course of differentiation. We uncovered that a number of Notch signaling responsive transcription factors (*Enhancer of split Complex*) are enriched in 4- and 8-cell cysts raising a possibility that Notch signaling regulates early steps of GC differentiation.

Two recent studies have generated similar ovary atlases (Rust et al. 2020, Jevitt et al. 2020). Each study had a distinct focus and approach (Supplemental Text). Our study and Rust *et al*. focused on early- and mid-stages of oogenesis. The main focus of Rust *et al.* study was the stem cell compartment, and they demonstrated that in specific environmental and genetic conditions the most posterior ECs can function as FSCs to give rise to FCs. Jevitt *et al.* sequenced the entire ovary and connected tissues, and characterized transcriptional trajectories of the entire follicle-cell population over the course of their development from stem cells to the oogenesis-to-ovulation transition. Moreover, they characterized the transcriptomes of the muscle sheath, hemocytes and oviduct. Overall, all three studies produced similar results, and majority of markers identified showed the expected expression in our dataset (Supplemental Figure 5, Supplemental Text). The exact cluster boundaries occasionally differed between the studies, for example, for EC subtypes. More focused studies will be needed to fully understand anatomical, morphological and functional diversity of ECs.

Altogether, the results of three independent scRNA-seq studies, including ours, are highly consistent despite technical differences in fly lines, sample preparation and analytical methods. They each present a unique dataset for further analyses. Our study provides a more granular sub-clustering of ovarian cell types, and precise cluster mapping to cell types and differentiation stages by direct visualization of mRNAs *in situ*. Thus, we deliver a comprehensive resource of gene expression profiles and markers for each cluster, and provide gene class annotations for transcriptional signatures and functional predictions.

Despite being one of the most extensively studied adult organs, our current work in the *Drosophila* ovary reveals higher cell type diversity than previously anticipated. These findings suggest that numerous, yet unidentified, cell subpopulations with distinct functions exist even in the most thoroughly studied organs. The ongoing Human Cell Atlas and similar projects in model organisms will start revealing this complexity, while focused studies will uncover the interplay between the subpopulations and their functions, finally allowing us to fully comprehend organ function in homeostasis and disease.

## Supporting information

Table 1 Markers

Table 2 Expression

Table 3 GLAD

Table 4 Predicted interaction

## Acknowledgements

The alpha-spectrin antibody developed by D. Branton and R. Dubreuil was obtained from the Developmental Studies Hybridoma Bank, created by the NICHD of the NIH and maintained at The University of Iowa, Department of Biology, Iowa City, IA 52242. We thank Drs. Katja Rust and Amy Reilein for help identifying follicle stem cells in stained germaria. We thank Drs. Katja Rust and Todd Nystul for sharing their results prior to publication. We thank Drs. Torsten Banisch, Sherilyn Grill, and Lionel Christiaen for critical comments on the manuscript. We would like to thank the Genome Technology Center (GTC) for expert library preparation and sequencing. GTC is a shared resource partially supported by the Cancer Center Support Grant P30CA016087 at the Laura and Isaac Perlmutter Cancer Center. Cell sorting technologies were provided by NYU Langone’s Cytometry and Cell Sorting Laboratory, which is supported in part by grant P30CA016087 from the National Institutes of Health/National Cancer Institute. S.G. was supported by Dean’s Undergraduate Research Fund Grant, R.L. was supported by the Simons Foundation, NIH R37HD41900 and was an HHMI investigator.

## Materials and Methods

### Fly husbandry

*w^1118^* flies were used for all stainings. *w^1118^; P[His2AV]62A* (Clarkson and Saint 1999) (BDSC #5941) flies crossed to *w^1118^* were used for all scRNA-seq experiments. Flies were raised on medium containing yeast, molasses and cornmeal, and kept at 25°C. Adult females were fattened on yeast for 24-48 hours prior to dissections.

### Ovary dissociation for scRNA-seq

For each sample, 30-40 pairs of adult ovaries were dissected one by one in ice-cold DPBS, no calcium, no magnesium (Thermo Fisher Scientific #14190136). Abdomens were removed using forceps and parts of intestine and abdominal cuticle were removed. Using forceps and a dissection needle, we cut off the anterior tip of the ovary and transferred to a new well.

#### Method A

For dissociation, anterior tips of ovaries were incubated in dissociation solution (0.5% Type I Collagenase (Thermo Fisher Scientific #17018029), 1% Trypsin 1:250 (Thermo Fisher Scientific #27250018) in DPBS) for 15 minutes with gentle rotation. The suspension was vigorously pipetted multiple times during the dissociation to enhance the dissociation efficiency. Enzymatic dissociation was stopped by adding Schneider cell culture media (Fisher # 21720-001) with 10% fetal bovine serum (Invitrogen #10082-147) (S-FBS). Starting from this step, all plastic materials – pipet tips, tubes, filters – were coated with S-FBS. Cell suspension was filtered through a custom-made ~40-micron cell strainer. The strainer was built by securing nylon mesh (Component Supply #U-CMN-40) in a cap of a 0.2 ml PCR tube and cutting the bottom of the tube and the cap (Slaidina et al. 2020). Upon filtering, dissociated cells were purified by fluorescence activated cell sorting (FACS) using a 100-micron nozzle on Sony SY3200 Cell Sorter. Cell suspension was centrifuged for 5 minutes at 250 rcf at RT, and resuspended in S-FBS.

#### Method B

For dissociation, anterior tips of ovaries we transferred to a 1.5 ml tube and were incubated in dissociation solution (0.25 mg/ml Type I Collagenase, 0.4 mg/ml elastase (Worthington Biochemical #LS002290) in Cell Dissociation Buffer (Thermo Fisher Scientific #13151014)) for 30 minutes with gentle rotation. The suspension was vigorously pipetted every 5 minutes during the dissociation to enhance the dissociation efficiency. Enzymatic dissociation was stopped by adding Schneider cell culture media with fetal bovine serum (S-FBS). Starting from this step, all plastic materials – pipet tips, tubes, filters – were coated with S-FBS. Cell suspension was filtered through a 70-micron filter (Miltenyi Biotec #130-095-823), and 30 micron filter (Miltenyi Biotec #130-041-407). Cell suspension was centrifuged for 5 minutes at 3500 rcf at 4^0^C. Cells were resuspended in 0.04% ultrapure BSA (Thermo Fisher Scientific #AM2616) in DBPS.

### Library preparation and sequencing

For scRNA-seq library preparation we used 10x Genomics Chromium Single Cell 3’ reagent Kits v2 for Method A (samples 1 and 2), and v3 for Method B (samples 3, 4 and 5) following the manufacturer's protocol. Sample 1 and 2 libraries were sequenced on paired-end 26/98 Illumina HiSeq 4000, and sample 3, 4, and 5 libraries on paired-end 28/91 NovaSeq 6000 runs.

### 10x Genomics data preprocessing

Per-read per-sample FASTQ files were generated using the Illumina bcl2fastq Conversion software (v2.20) to convert BCL base call files outputted by the sequencing instrument into the FASTQ format.

The 10X Genomics analysis software, Cell Ranger (v2.1.0 for samples 1 and 2, and v3.0.0 for samples 3,4 and 5), specifically the “cellranger count” pipeline, was used to process the FASTQ files in order to align reads to the *Drosophila melanogaster* reference genome (dm6) (Santos et al. 2015) and generate gene-barcode expression matrices. The output of multiple samples from the “cellranger count” pipeline were aggregated using the “cellranger aggr” pipeline of Cell Ranger, normalizing the combined output to the same sequencing depth and recomputing the gene-barcode matrices and expression analysis accordingly for the aggregated data.

### scRNA-seq data analysis

scRNA-seq datasets were integrated (aligned) using Seurat v3 (Stuart et al. 2019). We followed the Seurat v3 guidelines for identification of variable genes, dimensionality reduction and cell clustering. For step wise clustering, we selected cell groups from initial coarse clustering – GCs, follicle cells or somatic cells of germaria. The data was scaled, new PCAs were computed, and new UMAP plots were generated, followed by clustering with higher resolution parameters. To find markers, we used Wilcox statistical test built in Seurat 3 (Stuart et al. 2019). To select markers for visualization using HCR, we assessed their expression levels in cluster of interest and other clusters (Supplemental Figure S2). Importantly, we assured that GC markers were absent or expressed at very low levels in ECs or early FC lineages, and vice versa. Transcriptional signatures consisted of all positive cluster markers computed with 0.25 log fold change threshold. We used *pheatmap* package to generate heatmaps; gene expression levels were scaled by row. For all marker gene heatmaps, if a gene was assigned as a marker to multiple clusters, it was visualized in heatmaps only once, together with the markers of first cluster according to our cluster order. For example, a marker for 4-cc and 8-cc, was visualized together with 4-cc markers. We used the GLAD online tool to determine if the marker genes for each cluster fall into particular gene categories, and whether they are significantly enriched in each cluster’s transcriptional signature. For physical interaction analyses, we selected genes from following GLAD categories: GPCRs, Ion channels, Matrisome (ECM), Receptors, Secreted proteins, Transmembrane proteins, and Transporters (Hu et al. 2015).

### Monocle pseudotime analyses

We used Monocle 2 for pseudotime analyses of FSC differentiation (Qiu et al. 2017). We analyzed transcriptomes only from FSC/pre-FC, pre-stalk, stalk and polar cell cluster from samples 3, 4 and 5. These sample were processed in parallel on the same day, and thus, we were able to avoid any possible batch effects in our analyses.

### RNA in situ hybridization and immunofluorescence

All steps were done in using RNAse free reagents and supplies with gentle rotation, except for steps at 37°C. The protocol was adapted from Choi *et al*. (Choi et al. 2018). Custom designed probes, probe hybridization buffer, probe wash buffer and amplification buffer were procured from Molecular Instruments, Inc.

Specimens were fixed in 0.1% Tween (Tw), 4% Paraformaldehyde (PFA) (Electron Microscopy Sciences #15713) in DPBS for 20 minutes at RT, washed twice with 0.1% Tw in DPBS at room temperature (RT), dehydrated with sequential washes with 25%, 50%, 75% and 100% ethanol in DPBS on ice 5 minutes each. Samples were stored at −20°C at least overnight (up to 7 days). Samples were rehydrated with sequential washes with 100%, 75%, 50% and 25% ethanol in DPBS on ice, permeated for 2 hours in 1% Triton-X (Tx) DPBS at RT, post-fixed in 0.1% Tw, 4% PFA in DPBS for 20 minutes at RT, washed twice with 0.1% Tw in DPBS for 5 minutes on ice, washed with 0.1% Tw 0.5x DPBS 2.5xSSC (Thermo Fisher Scientific #AM9770) for 5 minutes on ice, washed twice with 5xSSCT (0.1% Tw in 5xSSC) for 5 minutes on ice, incubated in probe hybridization buffer for 5 minutes on ice, pre-hybridized in probe hybridization buffer for 30 minutes at 37°C, and hybridized in probe solution overnight (16 – 24 hours) at 37°C. Probe solution was prepared by adding probes to pre-warmed probe hybridization solution. Probe concentrations were determined empirically, and ranged 4-8 pmol of each probe in 1 ml. After hybridization, specimens were washed 4 times with probe wash buffer for 15 minutes each at 37°C, and twice with 5xSSCT for 5 minutes each at RT. Specimens were equilibrated in amplification buffer for 5 minutes at RT. Hairpin solutions were prepared by heating 30 pmol of each hairpin for 90 seconds at 95°C and cooling at RT in a the dark for 30 minutes, and subsequently adding the snap-cooled hairpins to 500 μl of amplification buffer at RT. Specimens were incubated in hairpin solution overnight (~16 hours) at RT, and washed multiple times with 5xSSCT – twice for 5 minutes, twice for 30 minutes and once for 5 minutes. If samples were not subjected to immunofluorescence staining, Dapi was added to the first 30 minute wash, and incubated in mounting media after completing all washes. For immunofluorescence samples were blocked in 0.1% Tw, 5% normal goat serum (NGS) (Jackson ImmunoResearch #005-000-121) in PBS at RT for 2 hours. Primary antibody was diluted in 0.1% Tw, 5% NGS in PBS and incubated for 2h at RT or 4 °C overnight. Subsequently, specimens were washed in 0.1% Tw in PBS times for 20 minutes in RT and in 0.1% Tw, 5% NGS in PBS twice for 30 minutes. Secondary antibody and Dapi (10 μg/ml for mounting in Vectashield, and 100 μg/ml for mounting in Slowfade Diamond) was diluted in 0.1% Tw, 5% NGS in PBS and incubated for 2h at RT or 4 °C overnight. Subsequently, specimens were washed in 0.1% Tw in PBS four times for 20 minutes in RT. Finally, specimens were equilibrated in Slowfade Diamond (Thermo Fisher Scientific #S36963) or Vectashield (Vector Labs Inc #H-1000) mounting medium at 4 °C overnight (or longer) and mounted in Slowfade Diamond or Vectashield mounting media. Primary antibody - anti-α-Spectrin (DSHB #3A9, concentrated) dilution 1:200. Secondary antibodies were procured from Jackson ImmunoResearch Laboratories, Inc.

### Imaging

Imaging was performed using Zeiss LSM 800 and Zeiss LSM 780 confocal microscopes using 40x oil NA 1.3 objectives.

## Data availability

The scRNA-seq data will be deposited in GEO.

## Notes

### Competing Interest Statement

The authors have declared no competing interest.

### Summary of Updates

The author list has been updated.

