## Supplemental Figures for "A single cell atlas reveals unanticipated cell type complexity in *Drosophila* ovaries"

A

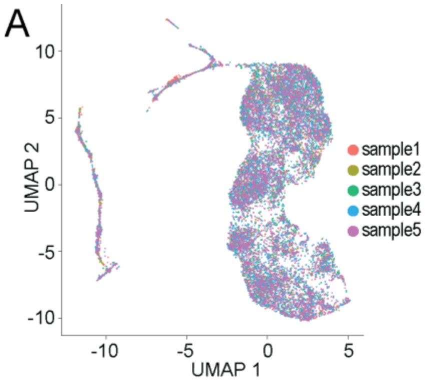


B

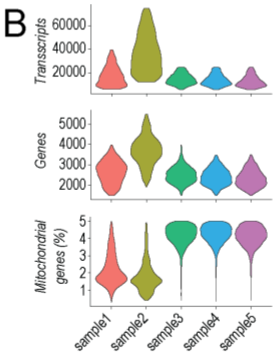


C

| | total cells | high quality cells | sequencing saturation | total genes detected | reads per cell |
|---|---|---|---|---|---|
| sample 1 | 1955 | 1450 | 78.0% | 11,850 | 182,576 |
| sample 2 | 1052 | 874 | 81.0% | 11,724 | 337,747 |
| sample 3 | 7192 | 3729 | 28.9% | 12,788 | 22,991 |
| sample 4 | 7794 | 3946 | 27.5% | 12,846 | 21,762 |
| sample 5 | 8559 | 5228 | 26.1% | 12,777 | 19,906 |
| total | 26552 | 15227 | | | |

D

| GSC/CB/ 2-cc | 4-cc | 8-cc | 16-cc 2a I | 16-cc 2a II | 16-cc 2ab | 16-cc 2b | 16-cc 3 | st 2 | |
|---|---|---|---|---|---|---|---|---|---|
| 3.0E-05 | 6.2E-06 | 4.5E-04 | 2.4E-04 | 4.2E-03 | 0.074 | 0.097 | 0.006 | 0.040 | Trans-membrane proteins |
| 2.9E-04 | 8.2E-08 | 0.010 | 0.034 | 0.048 | 0.032 | 0.228 | 0.108 | 0.162 | Cytoskeletal |
| 5.9E-03 | 8.8E-06 | 0.009 | 0.011 | 0.060 | 0.039 | 2.8E-04 | 1.6E-05 | 1.1E-13 | Metabolic vs2 |
| 0.007 | 0.002 | 0.012 | 0.007 | 0.004 | 0.133 | 0.149 | 0.138 | 0.127 | Secreted proteins |
| 4.0E-03 | 0.009 | 0.042 | 0.002 | 0.206 | 0.260 | 1.000 | 0.020 | 0.025 | Chaperone and heat shock proteins |
| 0.009 | 0.012 | 0.092 | 0.108 | 0.081 | 0.119 | 0.209 | 0.112 | 0.118 | Receptors |
| 0.058 | 0.002 | 0.015 | 0.007 | 0.001 | 0.120 | 0.155 | 0.006 | 0.016 | Transcription factor/DNA binding |
| 0.193 | 0.101 | 0.091 | 0.096 | 2.2E-04 | 0.031 | 0.008 | 0.163 | 0.043 | Major signaling pathways |
| 0.061 | 0.118 | 0.125 | 0.112 | 0.004 | 0.061 | 0.005 | 0.000 | 6.1E-11 | Mitochondrial |
| 0.021 | 0.021 | 0.151 | 0.139 | 0.058 | 0.153 | 0.064 | 0.184 | 2.7E-06 | Transporters |
| 0.274 | 0.223 | 1.000 | 0.230 | 0.236 | 1.000 | 0.213 | 0.109 | 0.001 | Ribosome |
| 0.024 | 0.054 | 0.151 | 0.120 | 0.048 | 0.073 | 0.142 | 0.072 | 0.006 | RNA-binding |

E

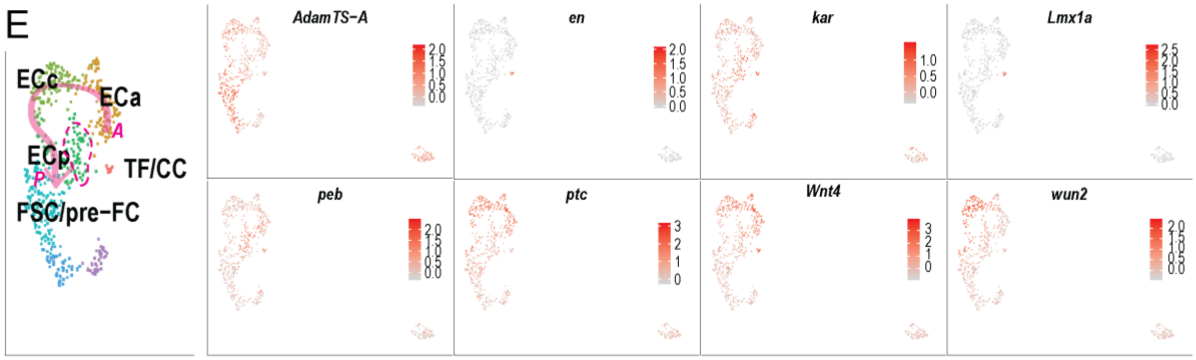


F

| TF/CC | aEC | cEC | pEC | FSC/ pre-FC | pre-stalk | stalk | polar | |
|---|---|---|---|---|---|---|---|---|
| 4.8E-04 | 4.1E-15 | 6.4E-08 | 0.243 | 0.370 | 0.001 | 8.3E-06 | 1.1E-06 | Major signaling pathways |
| 0.057 | 2.4E-05 | 0.007 | 0.160 | 0.118 | 0.019 | 0.060 | 0.005 | Secreted proteins |
| 3.1E-05 | 2.7E-04 | 0.166 | 0.315 | 0.284 | 0.057 | 0.015 | 0.272 | Chaperone and heat shock proteins |
| 0.006 | 2.9E-04 | 0.010 | 0.271 | 0.008 | 3.0E-06 | 2.4E-04 | 1.8E-08 | Matrisome |
| 0.002 | 0.050 | 5.1E-07 | 0.011 | 0.368 | 0.001 | 1.4E-05 | 1.9E-06 | Glycoproteins |
| 0.004 | 0.042 | 1.9E-04 | 6.9E-05 | 0.130 | 0.033 | 0.074 | 0.025 | Metabolic vs2 |
| 0.060 | 0.043 | 6.8E-04 | 0.250 | 0.304 | 0.103 | 0.025 | 0.141 | Receptors |
| 0.012 | 0.049 | 0.003 | 0.108 | 0.095 | 0.083 | 0.007 | 0.050 | Trans-membrane proteins |
| 0.105 | 0.264 | 0.004 | 0.032 | 0.335 | 0.241 | 0.171 | 1.000 | Autophagy-related |
| 0.148 | 0.021 | 0.009 | 0.351 | 0.370 | 0.069 | 0.173 | 0.142 | Kinases |
| 0.062 | 0.060 | 0.029 | 4.2E-04 | 0.092 | 4.0E-04 | 3.6E-03 | 0.060 | Mitochondrial |
| 0.023 | 0.079 | 0.028 | 0.319 | 4.2E-05 | 1.4E-06 | 8.4E-06 | 3.2E-03 | Cytoskeletal |
| 0.034 | 0.011 | 0.066 | 0.030 | 0.127 | 0.010 | 0.025 | 0.030 | Transcription factor/DNA binding |

**Supplemental Figure 1.**

A - UMAP plot of entire dataset visualizing integration of 5 *[Continues on the next page]*

*[Continued from the previous page]* samples. Each dot represents a transcriptome of a single cell, and is color-coded according to sample. B - Violin plots visualizing number of transcripts detected, genes detected and fraction of mitochondrial reads in each sample. C - A table showing experiment statistics - total number of cells, number of high-quality cells, sequencing saturation, total genes detected and reads per cell. D - A table showing p-values of gene class enrichment in GC cluster markers. Darker purple shading indicates lower p-values. E - Feature plots visualizing select marker gene expression in somatic cells of germarium. Left panel - fragment of UMAP plot from Figure 3A. Right panel - visualization of *AdamTS-A*, *en*, *kar*, *Lmx1a*, *peb*, *ptc*, *Wnt4* and *wun2* expression in somatic cells of germarium. Each dot represents a cell, and is color-coded according to gene expression levels as indicated in the scale. F - A table showing p-values of gene class enrichment in somatic cell of germarium cluster markers. Darker purple shading indicates lower p-values.

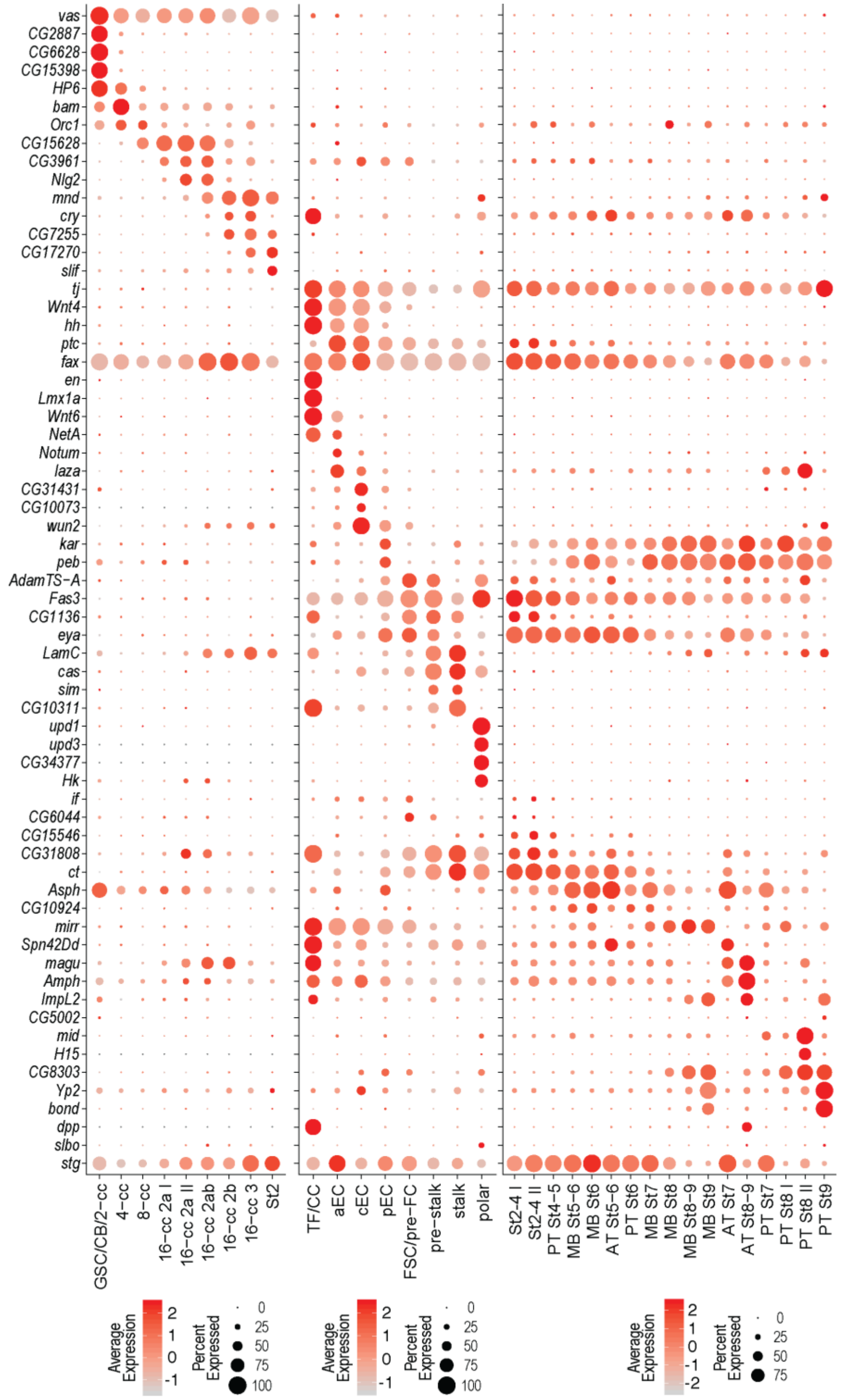


**Supplemental Figure 2.**
Dot plot visualizing expression of all marker genes used in our study. Dot diameter represents the fraction of cells expressing each gene in each cluster, as shown in scale. Color intensity represents the average normalized expression level.

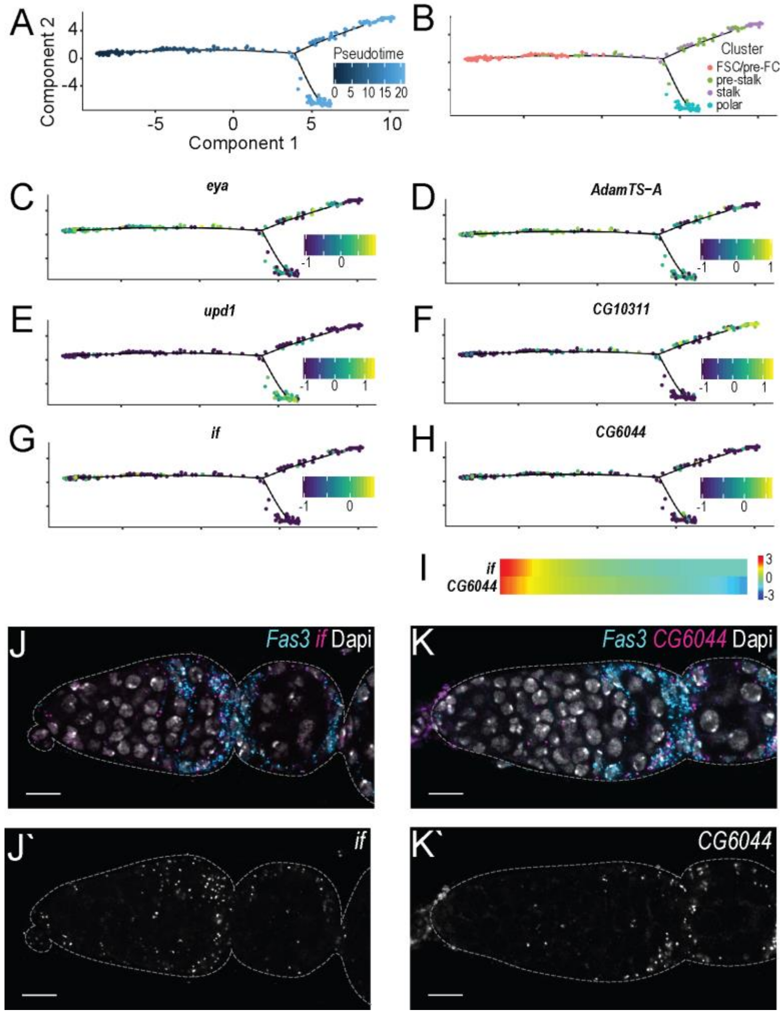


**Supplemental Figure 3.**

A-H - pseudotime plots visualizing FSC differentiation trajectory. Each dot represents a cell and is color-coded according to: A - 'pseudotime' as shown in scale; B - cluster identity as shown in legend; C - *eya* expression as shown in scale; D - *AdamTS-A* expression as shown in scale; E - *upd1* expression as shown in scale; F - *CG10311* expression as shown in scale; G - *if* expression as shown in scale; H - *CG6044* expression as shown in scale. I - heatmap visualizing expression of two putative FSC markers *if* and *CG6044*. Red indicates high and blue low relative expression as shown in scale. Early timepoints are on the left and late on the right. J, K - Marker gene mRNA *in situ* hybridization using HCR. *Fas3* (cyan) is expressed in FCs, Dapi (white) labels cell nuclei. FCs have characteristic elongated nuclei, while GCs have rounded nuclei. Scale bars, 10 µm. J - *if* (magenta, white) is expressed in FSCs, pre-FCs, and sparsely and at low levels in ECs. K - *CG6044* (magenta, white) is expressed predominantly in pre-FCs, with lower levels in FSCs.

A

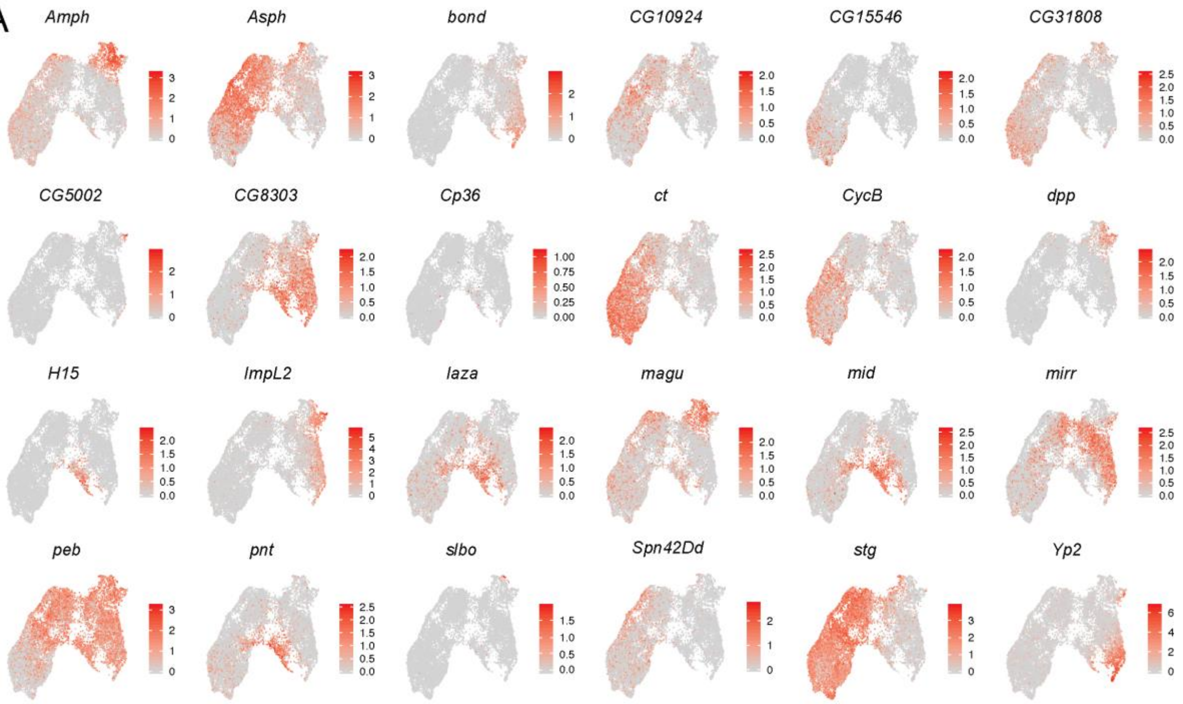


B

| St2-4 I | St2-4 II | PT St4-5 | MB St5-6 | MB St6 | MB St7 | MB St8 | MB St8-9 | Mb St9 | |
|---|---|---|---|---|---|---|---|---|---|
| 4.3E-07 | 0.001 | 4.0E-04 | 0.001 | 0.093 | 0.232 | 0.283 | 0.222 | 0.009 | Transcription factor/DNA binding |
| 4.3E-04 | 2.2E-09 | 0.017 | 0.063 | 0.001 | 0.309 | 0.366 | 0.375 | 0.195 | Cytoskeletal |
| 0.003 | 0.000 | 0.029 | 0.060 | 0.025 | 0.239 | 0.060 | 5.1E-06 | 6.2E-09 | Metabolic vs2 |
| 0.004 | 0.001 | 4.3E-04 | 0.079 | 0.085 | 0.104 | 0.116 | 0.163 | 0.002 | Trans-membrane proteins |
| 1.9E-05 | 0.008 | 0.002 | 0.042 | 0.362 | 0.279 | 1.000 | 1.000 | 1.000 | Major signaling pathways |
| 0.002 | 0.044 | 0.105 | 0.348 | 1.000 | 1.000 | 0.207 | 0.110 | 0.081 | Matrisome |
| 0.015 | 0.001 | 0.071 | 0.076 | 1.000 | 0.288 | 0.015 | 4.0E-04 | 7.8E-06 | Metabolic |
| 0.078 | 0.048 | 0.130 | 0.247 | 1.000 | 0.380 | 0.274 | 1.9E-05 | 0.000 | Mitochondrial |
| 0.181 | 0.070 | 0.200 | 0.341 | 1.000 | 0.361 | 0.079 | 3.7E-04 | 0.001 | Human Disease |
| 1.000 | 1.000 | 1.000 | 1.000 | 1.000 | 1.000 | 0.024 | 0.036 | 1.6E-04 | Chaperone and heat shock proteins |
| 0.036 | 0.035 | 0.066 | 1.000 | 1.000 | 1.000 | 1.000 | 0.037 | 0.002 | Transporters |
| 0.165 | 0.139 | 0.147 | 0.275 | 0.257 | 0.040 | 0.244 | 0.272 | 0.007 | Secreted proteins |
| 0.167 | 0.198 | 0.206 | 0.372 | 0.357 | 0.059 | 0.352 | 0.043 | 0.017 | Glycoproteins |
| 0.257 | 0.182 | 0.358 | 1.000 | 0.118 | 0.181 | 0.253 | 0.063 | 0.023 | Autophagy-related |

C

| AT St5-6 | AT St7 | AT St8-9 | |
|---|---|---|---|
| 2.7E-05 | 0.003 | 0.011 | Cytoskeletal |
| 0.006 | 8.3E-04 | 3.2E-05 | Matrisome |
| 0.001 | 0.152 | 0.146 | Transcription factor/DNA binding |
| 0.009 | 0.028 | 0.152 | Major signaling pathways |
| 0.040 | 0.008 | 4.0E-04 | Glycoproteins |
| 0.105 | 0.067 | 6.3E-07 | Metabolic vs2 |
| 0.035 | 0.135 | 1.6E-04 | Metabolic |
| 1.000 | 0.276 | 4.8E-04 | Transporters |
| 0.196 | 0.186 | 0.002 | Mitochondrial |

D

| PT St6 | PT St7 | PT St8 I | PT St8 II | PT St9 | |
|---|---|---|---|---|---|
| 4.6E-05 | 0.366 | 0.014 | 1.000 | 0.153 | Cytoskeletal |
| 0.024 | 0.024 | 0.001 | 0.122 | 8.5E-05 | Trans-membrane proteins |
| 0.351 | 0.199 | 0.004 | 0.094 | 0.014 | Chaperone and heat shock proteins |
| 0.174 | 0.207 | 0.007 | 0.260 | 0.002 | Matrisome |
| 0.172 | 0.119 | 0.008 | 0.004 | 1.3E-07 | Glycoproteins |
| 0.149 | 0.097 | 0.044 | 2.6E-12 | 9.8E-10 | Metabolic |
| 0.125 | 0.114 | 0.101 | 5.4E-12 | 2.7E-10 | Metabolic vs2 |
| 0.049 | 0.151 | 0.085 | 9.0E-08 | 3.1E-09 | Mitochondrial |
| 0.181 | 0.079 | 0.178 | 7.1E-06 | 0.001 | Human Disease |
| 1.000 | 0.253 | 1.000 | 1.4E-04 | 0.015 | Autophagy-related |
| 0.162 | 0.185 | 0.121 | 0.107 | 3.3E-04 | Transcription factor/DNA binding |
| 0.213 | 0.106 | 0.018 | 0.137 | 0.001 | Secreted proteins |

**Supplemental Figure 4.**

A - Feature plots visualizing previously known and newly identified FC marker gene expression in alphabetical order. Each dot represents a cell and is color-coded according to gene expression levels, as indicated in the scale. B-D - Tables showing p-values of gene class enrichment in FC cluster markers. Darker purple shading indicates lower p-values. B - early stage and MBFCs, C - ATFCs, D - PTFCs.

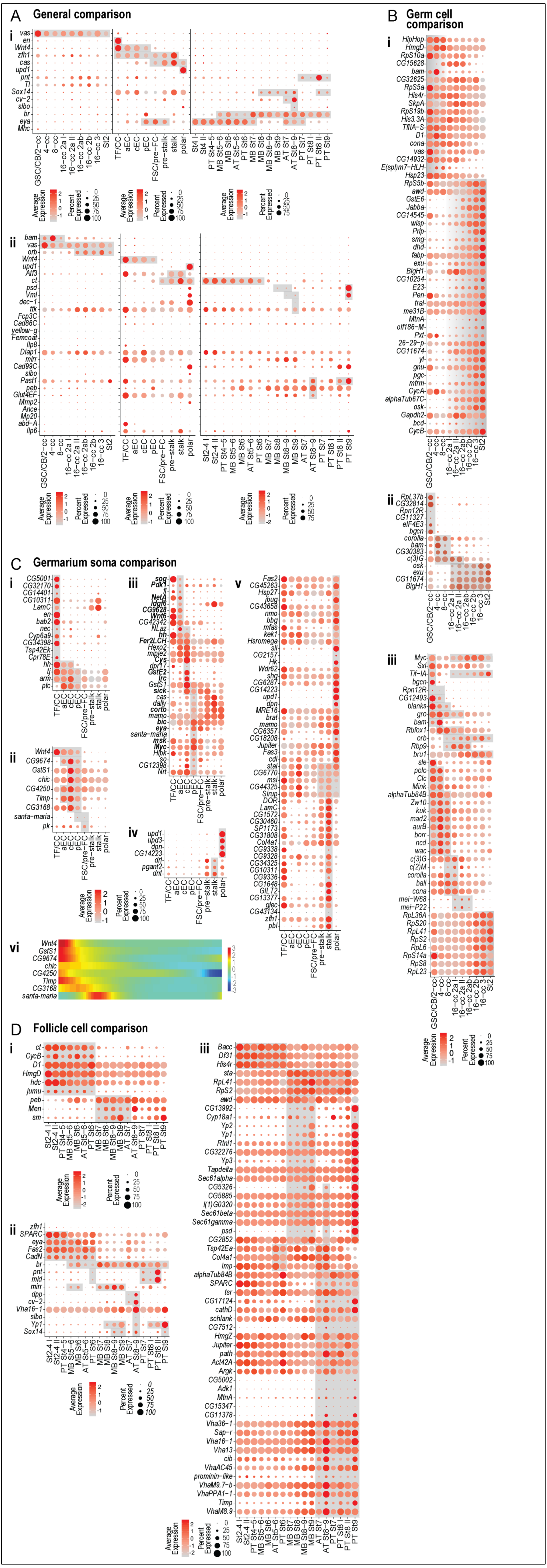

A General comparison
B Germ cell comparison
C Germarium soma comparison
D Follicle cell comparison
Average Expression
Percent Expressed

**Supplemental Figure 5.**
A-D - Dot plots visualizing expression of marker genes from Jevitt *et al.* and Rust *et al.* in our dataset. Dot diameter represents the fraction of cells expressing each gene in each cluster, as shown in scale. Color intensity represents the average normalized expression level. A - general markers from Rust et al. (i) and Jevitt et al. (ii). B - GC markers from Jevitt *et al*. pseudotime analyses (i), Rust *et al.* clustering (ii) and pseudotime analyses (iii). C - germarium soma markers. TF and CC markers (i), FSC markers (ii), EC markers (iii), polar and stalk cell markers (iv) from Rust *et al*., polar and stalk cell markers from Jevitt *et al*. pseudotime analyses (v). C(vi) - heatmap visualizing expression of putative FSC markers from Rust *et al*. in our FSC differentiation pseudotime analyses. Red indicates high and blue low expression as shown in scale. Early timepoints are on the left and late on the right. D - follicle cell markers. Mitotic and endocycling FC markers from Jevitt *et al*. (i), all FC markers from Rust *et al*. (ii), and MB and terminal FC enriched genes from Jevitt *et al*. pseudotime analyses (iii).
